# Distinct matrix viscoelasticity in bone fracture hematoma steers macrophage polarization

**DOI:** 10.1101/2025.11.14.688455

**Authors:** Raphael S. Knecht, Duncan M. Morgan, Matthias R. Kollert, Anne Noom, Christian H. Bucher, David J. Mooney, Katharina Schmidt-Bleek, Georg N. Duda

**Author notes:** These authors contributed equally as second authors (Duncan M. Morgan, Matthias R. Kollert).

## Abstract

Physical properties of the extracellular matrix (ECM) are key regulators of cellular behavior. Following injury, the formation of a hematoma establishes a provisional niche that initiates and regulates healing responses. However, the influence of hematoma viscoelastic properties on immune cell behavior remains poorly understood. Here, we show that distinct ECM viscoelastic properties of the maturing fracture hematoma steer macrophage polarization from pro-inflammatory to pro-regenerative characteristics. Tissue analyses of *ex vivo* human samples revealed that hematoma viscoelastic properties change with ECM remodeling during healing progression, with the late-phase stress relaxation time constant, τ2, increasing significantly with days post-injury. Using alginate hydrogels in 3D culture, we engineered extracellular microenvironments with tunable τ₂ but constant stiffness to study their role in macrophage polarization. Our data demonstrate that ECM τ₂ properties guide macrophage phenotype, characterized by high τ₂ promoting pro-inflammatory activation, while low τ₂ supported anti-inflammatory phenotypes. This regulation of macrophage polarization by ECM stress relaxation properties persists even under toll-like receptor-coactivation. Single-cell RNA sequencing revealed distinct transcriptional programs associated with different ECM τ₂ values, with many of the differentially expressed genes related to metabolic processes. The transcriptomic profiles of macrophages primed by different ECM τ₂ aligned with *in vivo* healing trajectories, with the gene signature score of the low ECM τ2 decreasing over time. Our findings uncover the immune-regulatory function of specific hematoma stress relaxation properties associated with healing progress after injury, and suggest τ₂ as potential mechanobiological target to be leveraged in novel biomaterials-based regenerative therapies.

## Introduction

Following injury, the formation of a hematoma marks the onset of a cascade of healing processes. Beyond its hemostatic function, the hematoma serves as a dynamic cell niche that coordinates the complex interplay between immune responses, cellular recruitment, and tissue remodeling. In bone fracture healing, this initial phase is critical for enabling successful tissue regeneration^1^. The early progression of fracture healing is characterized by dynamic changes within the hematoma, including the transition from an initial pro-inflammatory phase to a subsequent anti-inflammatory phase, which enables scar-free tissue healing^2–4^. One key process of this transition is macrophage polarization, where pro-inflammatory macrophages (M1-like) shift toward anti-inflammatory phenotypes (M2-like). Pro-inflammatory macrophages clear cellular and tissue debris and secrete inflammatory cytokines that recruit additional immune cells and mesenchymal stem cells (MSCs)^5^. In contrast, anti-inflammatory macrophages release growth factors, such as vascular endothelial growth factor (VEGF), which promote the formation of new blood vessels, a crucial step for restoring oxygen and nutrient supply to the injury site and enabling further progression of the healing cascade^5–7^. Dysregulation of the transition from pro-to anti-inflammatory phases impairs bone regeneration, such as non-union or delayed healing, underscoring the clinical importance of this tightly regulated process^8–10^.

The extracellular matrix (ECM) in fracture hematoma also undergoes dramatic change during the proceeding healing. This ECM change is characterized by structural reorganization, progressing from an initial fibrin clot to a mature network of matrix fibers^11–13^. As the mechanical properties of ECMs are governed by their matrix structures^14^, the substantial matrix remodeling during hematoma maturation would be expected to influence the viscoelastic properties of the ECM. ECM viscoelastic properties, including elastic modulus and time-dependent stress relaxation, are inherent to biological tissues^15^. The elastic modulus of the ECM is an important regulator of macrophage polarization, with stiffer substrates promoting pro-inflammatory^16^ and softer substrates favoring anti-inflammatory phenotypes^17, 18^. Although the current understanding is limited to predominantly two-dimensional culture systems, failing to recapitulate the three-dimensionality of most extracellular microenvironments. Further, an almost exclusive focus on elastic properties neglects the viscoelastic characteristics of biological ECMs^19^. The stress relaxation properties of the ECM kinetics influence osteogenic differentiation of mesenchymal stromal cells^19^, fate decision of monocytes^20^, and function of T-cells^21^. However, it remains unclear how ECM viscoelastic properties changes during hematoma maturation, and whether associated changes in ECM mechanical cues might influence macrophage polarization within the context of bone healing after fracture.

We postulated that the evolving viscoelastic properties of fracture hematoma, driven by ECM maturation and reorganization, serve as critical biophysical cues that steer macrophage polarization during the initial phases of bone healing. To test this, we first characterized the viscoelastic properties of *ex vivo* human fracture hematoma samples at different stages of healing. Using these data, we used ionically crosslinked alginate hydrogels to mimic the ECM viscoelastic properties of specific stages of hematoma maturation in 3D culture *in vitro*. Human monocyte-derived macrophages were encapsulated in hydrogels and characterized via flow cytometry, cytokine profiling, and single-cell RNA sequencing. Finally, we compared the RNA-seq profiles of our *in vitro* data on macrophages primed by ECM mechanics with those of maturing fracture hematoma from an *in vivo* murine bone fracture model.

Our results reveal that the ECM viscoelastic properties change with the maturation of fracture hematoma, with specifically the late-phase stress relaxation (τ_2_) increasing over time. We show that the differences in ECM τ_2_, which were characterized in ex vivo tissue samples, can be sensed by macrophages and steer their polarization. Specifically, faster late-phase stress relaxation (low τ_2_) promotes pro-inflammatory polarization, while slower late-phase stress relaxation (high τ_2_) favors anti-inflammatory phenotypes. The modulation of macrophage polarization by low and high τ_2_ was independent of integrin-mediated adhesion or coactivation of toll-like receptors (TLRs). Our findings establish ECM viscoelastic properties as an extracellular physical regulator of macrophage polarization, and provide new insights for developing biomaterials to direct immune responses in therapies targeting tissue regeneration.

## Result

### Fracture hematoma viscoelastic properties correlate with days post injury

We first investigated whether the maturing and reorganizing ECM of the fracture hematoma undergoes changes in its viscoelastic properties during the early phase of healing (**Figure 1a**). To test this, we collected fracture hematoma samples from patients undergoing orthopedic surgery and performed uniaxial compression tests to characterize their viscoelastic properties (**Figure 1b**). Stress relaxation curves were fitted using both one-and two-element Maxwell models **(Figure 1c)**. Stress relaxation curves and fits from all fracture hematoma samples can be found in **Supplementary Figure 1**.The two-element model consistently yielded superior fits across all samples, as reflected by higher coefficients of determination (R²), lower root mean square error (RMSE), and strong support from information criteria **(Figure 1d,e** and **Supplementary Table 1)**. Likelihood ratio tests further confirmed that the improvement from one to two elements was highly significant (χ², *p* < 0.001, **Supplementary Table 2**). While extending the model to include additional Maxwell elements led to higher R² values and statistically significant improvements, these came at the expense of parameter stability and yielded biologically unlikely values. Importantly, the RMSE of the two-element model was already lower than the estimated level of experimental noise (**Supplementary Figure 2d**). Therefore, to preserve interpretability and avoid overfitting, we selected the two-element model as the optimal representation of stress relaxation behavior in human fracture hematoma. These results suggest the involvement of at least two viscoelastic processes: a rapid phase, potentially reflecting the rupture of weak molecular bonds, and a slower phase likely driven by ECM polymer rearrangement^15, 22–24^. The two-element Maxwell model effectively captures this biphasic behavior through its distinct time constants, the early-phase stress relaxation constant, τ₁, and the late-phase stress relaxation constant, τ₂.

**Figure 1:**
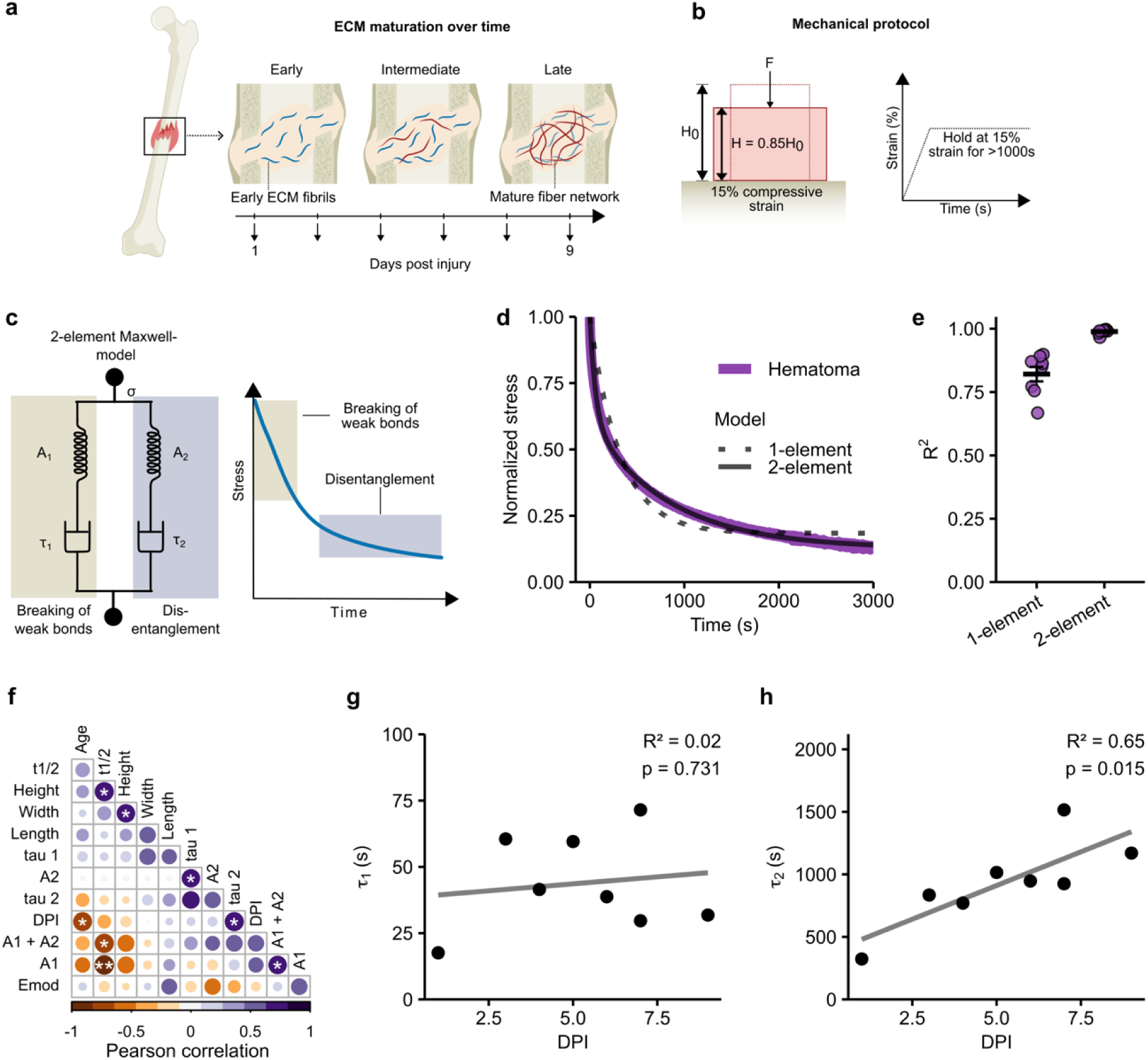
ECM viscoelastic properties in fracture hematoma correlate with healing progression. a) Schematic representation of the bone healing process. Early fracture hematoma is characterized by a loosely organized extracellular matrix (ECM), which progressively matures over time into a more structured and entangled ECM network. b) Experimental protocol for uniaxial compression testing of human fracture hematoma to assess viscoelastic properties. c) Stress-relaxation behavior was modeled using a two-element Maxwell model to capture both early (e.g., weak bond rupture) and late (e.g., ECM polymer disentanglement) relaxation processes. d) Representative stress-relaxation curve showing fits of the one-element and two-element Maxwell models. e) Goodness-of-fit comparison: the two-element model achieved significantly higher R² values (0.99 ± 0.01) than the one-element model (0.82 ± 0.08). Data are mean ± SEM from n = 8 biologically independent samples. f) Pearson correlation matrix of mechanical parameters derived from the model. Height, width, and length indicate sample dimensions; age denotes patient age; Emod is the elastic modulus; A₁ and A₂ are the relaxation amplitudes of each model branch, with A₁ + A₂ representing total relaxation; DPI indicates days post injury; τ₁ and τ₂ are the characteristic time constants of the early and late relaxation processes, respectively. g–h) Linear regression analyses of τ₁ (g) and τ₂ (h) with respect to days post injury.

To explore how these mechanical parameters change with time post-injury, we analyzed correlations between model-derived constants and days post injury (**Figure 1f** and **Supplementary Figure 3**). While stress relaxation half-time (t_1/2_) and τ_1_ showed no significant association with days post injury, τ_2_ displayed a significant positive correlation. Linear regression was consistent with this finding: τ_1_ remained unchanged over time **(Figure 1g)**, whereas τ_2_ increased significantly with days post injury (p = 0.015) (**Figure 1h**), suggesting a progressive reorganization of the ECM with healing is mainly affecting τ_2_.

### Tunable stress relaxation via ionic crosslinking and molecular weight of alginate hydrogels enables independent modulation of τ₂ without altering τ₁

To decouple the effects of distinct stress relaxation components on macrophage polarization, we used an *in vitro* hydrogel system that allowed independent tuning of the τ_2_, while maintaining a constant τ_1_ (**Figure 2a**). We selected ionically crosslinked alginate hydrogels for their versatility in mechanical tuning. We hypothesized that hydrogels formed from low molecular weight alginate resulted in matrices with low τ_2_ value due to fewer polymer entanglements. Conversely, using higher molecular weight alginate yielded more viscous matrices with high τ_2_ values, due to increased chain entanglement.

**Figure 2:**
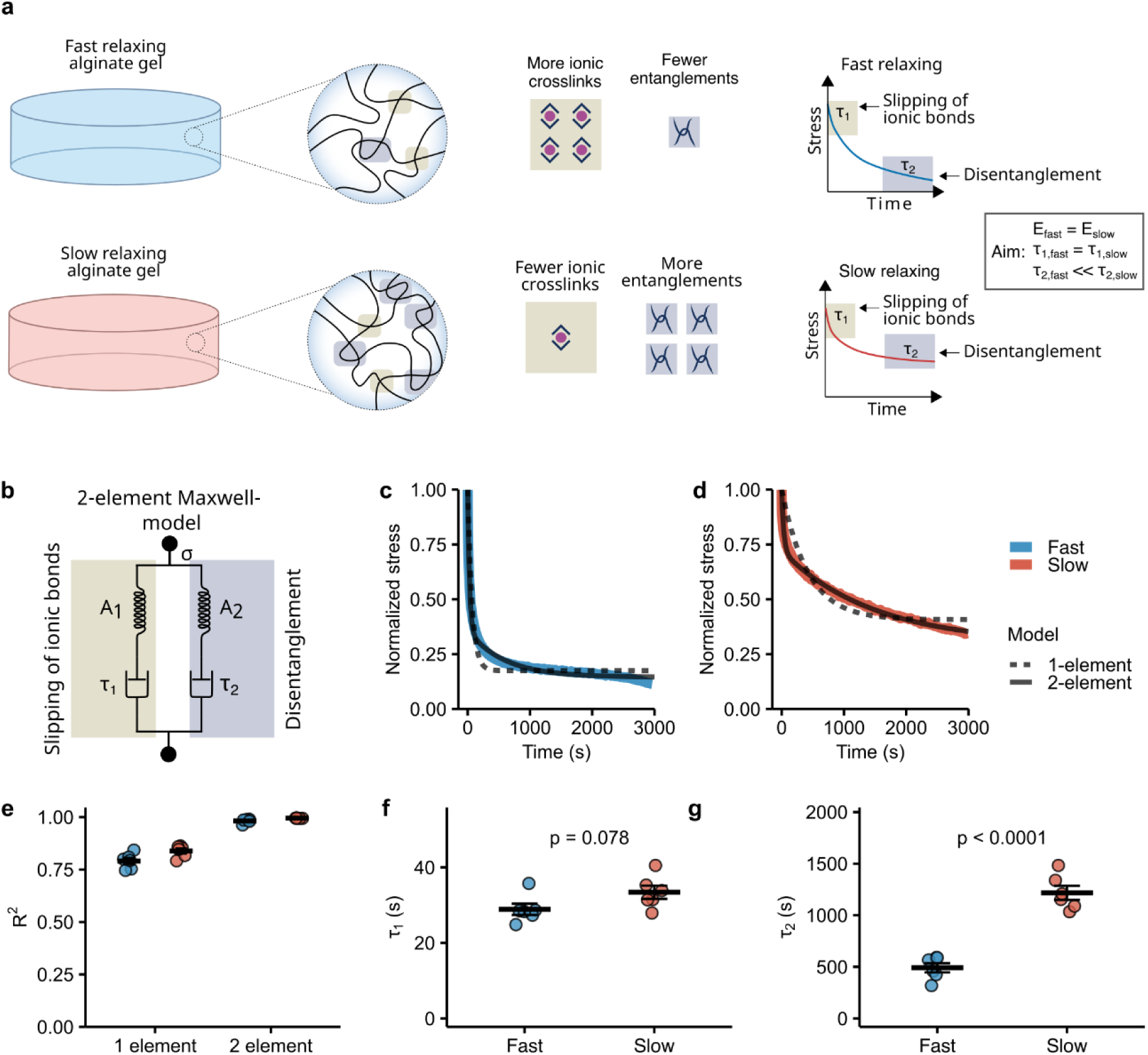
Independent modulation of ECM τ_1_ and τ_2_ using alginate hydrogels. a) Left: To modulate relaxation kinetics while maintaining a constant elastic modulus, fast relaxing hydrogels were formulated with low molecular weight alginate (reduced entanglement) and higher calcium concentrations (increased ionic crosslinking). In contrast, slow relaxing hydrogels used high molecular weight alginate (enhanced entanglement) and low calcium concentrations (fewer ionic crosslinks). Right: Early stress relaxation is dominated by ionic bond slippage, while late-phase relaxation is controlled by polymer chain disentanglement; resulting in distinct relaxation kinetics between fast and slow relaxing hydrogels. b) Stress-relaxation behavior was modeled using a two-element Maxwell model to capture both early (slipping of ionic bonds) and late (polymer disentanglement) relaxation processes. c–d) Representative normalized stress-relaxation curves for c) fast relaxing and d) slow relaxing alginate hydrogels, overlaid with curve fits from one-and two-element Maxwell models. e) Goodness-of-fit comparison: The two-element model yielded superior R² values (0.98 ± 0.01 for fast and 0.99 ± 0.00 for slow) compared to the one-element model (0.79 ± 0.04 for fast and 0.84 ± 0.03 for slow). f) τ₁ time constant (early component) and g) τ₂ time constant (late component) from the two-element Maxwell model. Data in e-g are mean ± SEM. from n = 5 independent alginate gels. P values were determined using two-tailed unpaired t-test.

Stress relaxation was measured after 15% compression and fitted using one-and two-element Maxwell models (**Figure 2b**). In the two-element model, τ_1_ captures rapid relaxation from ionic bond slippage, while τ_2_ reflects delayed relaxation driven by polymer chain mobility and disentanglement^25^. Consistent with the expected biphasic relaxation profiles, the two-element Maxwell model provided better fits for both fast and slow relaxing hydrogels with low and high τ_2_ respectively (**Figure 2c,d**). R² values were markedly higher for the two-element model (fast: 0.98 ± 0.01; slow: 0.99 ± 0.00) than for the one-element model (fast: 0.79 ± 0.04; slow: 0.84 ± 0.03) (**Figure 2e**). Model comparison using AIC/BIC and F-tests robustly favored the two-element Maxwell model across all conditions (ΔAIC/BIC > 750,000; p < 0.001). Importantly, low and high τ_2_ gels recapitulated the lower and upper τ_2_ values observed across the measured fracture hematoma healing time period (days 1-9).

Importantly, this approach yielded two distinct alginate gels with only a modest difference in τ₁ (p = 0.078) (**Figure 2f**), yet a significant difference in τ₂ (p < 0.0001) (**Figure 2g**). Fast relaxing hydrogels exhibited τ₂ = 491 ± 110s, whereas slow relaxing gels showed τ₂ = 1218 ± 167s. These results demonstrate that dual modulation of network formation (ionic and polymer network architecture) enables selective tuning of τ₂, whereas changes in τ_1_ are minimal. This mechanical decoupling provides a platform to investigate how distinct modes of stress relaxation, particularly slow acting mechanisms such as polymer chain disentanglement, influence macrophage polarization.

### ECM τ₂ differentially regulates surface marker expression and cytokine secretion of macrophages

Mechanical cues of the extracellular microenvironment regulate cell fate decisions in immune cells^20, 21, 26^. Here, we hypothesized that the changes of ECM τ_2,_ which we characterized in human *ex vivo* hematoma during the inflammatory phase of fracture healing, represent biophysical cues that modulate macrophage polarization. To test our hypothesis, we isolated human monocytes and differentiated them into macrophages using granulocyte-macrophage colony-stimulating factor (GM-CSF) over 4 days, then encapsulated these cells for 48 hours in alginate hydrogels with distinct τ_2_ (Figure 3a). During 4 days of GM-CSF stimulation, monocytes upregulated CD11b, CD86, CD206, and HLA-DR (**Supplementary Figure 4**). Macrophages were positive for CD45+CD11b+ and HLA-DR+ before encapsulation (**Supplementary Figure 5**).

**Figure 3:**
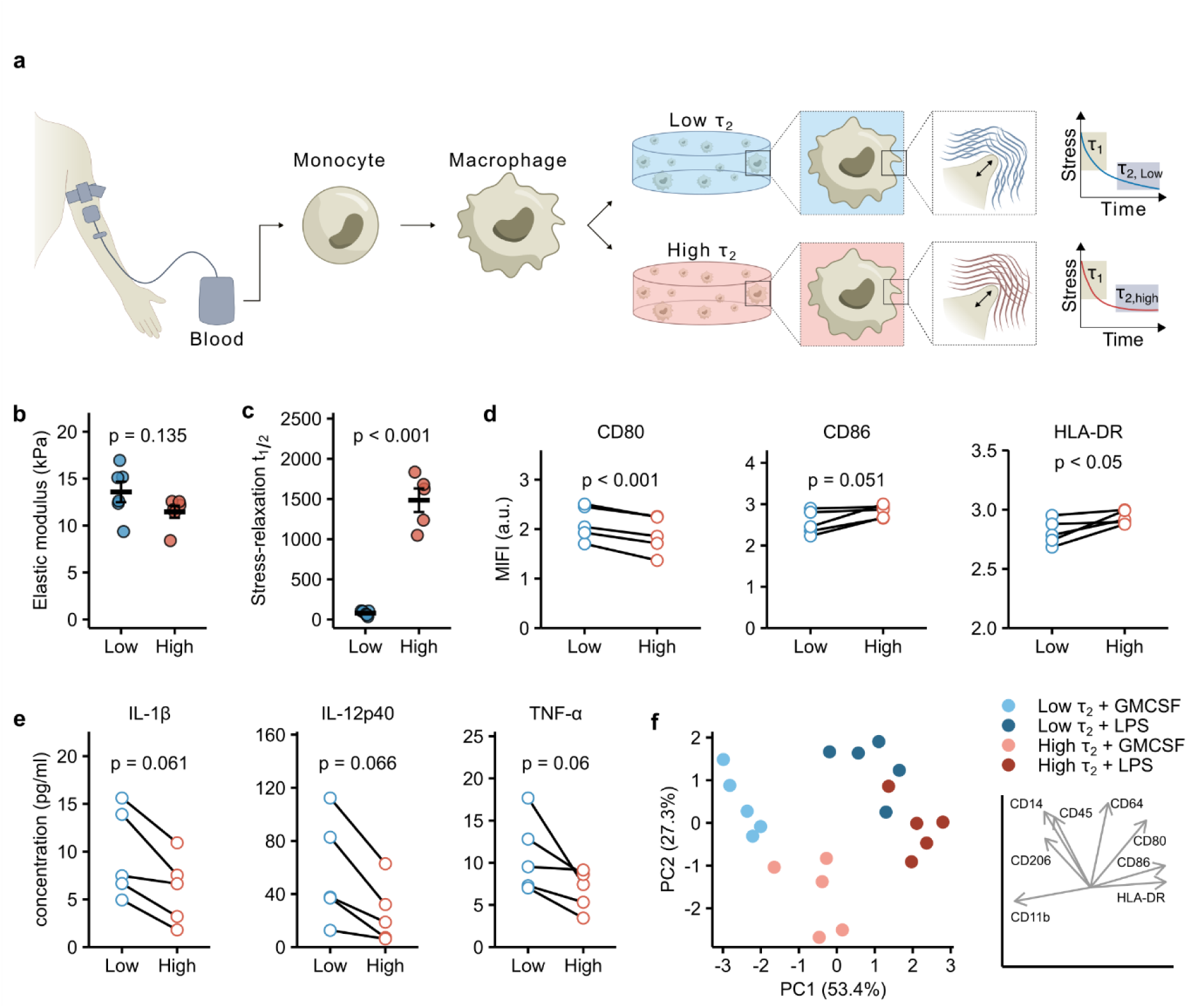
τ₂ differentially regulates macrophage surface marker expression and cytokine secretion. a) Experimental design: Human peripheral blood monocytes were isolated and differentiated into macrophages prior to encapsulation in alginate hydrogels with either low or high τ_2_. b) Young’s modulus of low and high τ_2_ alginate gels, measured at a strain rate of 1 mm/min between 8–12% strain. c) Stress relaxation half-time (t₁/₂) at 15% strain. d) Median logicle-transformed fluorescence intensity (MlFI) of CD80, CD86, and HLA-DR following 48 h encapsulation of macrophages in either low or high τ_2_ alginate hydrogels. e) Concentrations of secreted IL-1β, IL-12p40, and TNF-α in culture supernatant after 48 h encapsulation. f) Principal component analysis (PCA) of surface marker expression reveals consistent modulation by τ_2_ in GM-CSF and LPS-stimulated macrophages. Plot on the right depicts loadings of the first two principle components. Each data point represents an independent donor (n = 5). P-values determined by two-tailed unpaired t-test; n =5 for b) and c) and two-tailed paired t-test; n=5 for d) and e).

To isolate the effects of different ECM stress relaxation properties, we configured hydrogels with high or low ECM τ_2_ but constant stiffness. For this purpose, we adjusted calcium crosslinker concentrations and molecular weight (M_w_) of alginate chains, using 15 mM Ca^2+^ with high M_w_ alginate to form high τ_2_ hydrogels and 27mM Ca^2+^ with low M_w_ alginate to form low τ_2_ hydrogels, compensating for fewer network entanglements due to shorter alginate chains. This yielded alginate hydrogels with comparable initial stiffness (low τ_2_: 13.6 ± 2.7 kPa, high τ_2_: 11.5 ± 1.6 kPa, p = 0.135) (**Figure 3b**). Mesh size of alginate hydrogels is expected to stay constant with varying calcium concentration^27^ and molecular diffusion properties was comparable between conditions (**Supplementary Figure 6**). The τ_2_ values from the alginate hydrogels recapitulate the τ_2_ range of human fracture hematoma (τ_2=_ 323–1515 s) during the initial healing phase. Stress relaxation half-times (t_1/2_) were also distinct between the alginate gels (t_1/2, low_ = 55s, and t_1/2, high_ = 1587s) (**Figure 3c**). Notably, our two-element Maxwell model analysis revealed that the relative contributions of τ_1_ versus τ_2_ to the total amount of stress relaxation differ slightly between formulations: τ_2_ accounting for 29% of total stress relaxation in high τ_2_ gels compared to 38% in low τ_2_ gels (**Supplementary Table 3**). Flow cytometry analysis revealed that macrophages in low τ_2_ gels exhibited significantly elevated expression of the pro-inflammatory marker CD80 compared to high τ_2_ gels (p<0.001) **(Figure 3d**). Conversely, macrophages in high τ_2_ environments showed trends toward higher expression of CD86 (p = 0.051) and HLA-DR (p < 0.05), suggesting a shift toward regulatory phenotypes^28^. A full panel of measured surface markers is provided in **Supplementary Figure 7**. Cytokine secretion analysis supported these polarization differences, with macrophages in low τ_2_ gels showing a trend toward higher concentrations of pro-inflammatory mediators IL-1β (p = 0.059), IL-12p40 (p = 0.061) and TNF-α (p = 0.066) (**Figure 3e**). A full panel of measured cytokines is provided in **Supplementary Figure 8**.

To test whether the mechanical microenvironment modulates macrophage polarization also under pro-inflammatory conditions mimicking post-fracture inflammation, we co-stimulated macrophages with LPS and IFN-γ. LPS activates Toll-like receptor 4, the same receptor engaged by damage-associated molecular patterns (DAMPs) released during fracture^29, 30^. Under these conditions, macrophages upregulated the markers CD80, CD86, and HLA-DR in both low and high τ_2_ gels (**Supplementary Figure 11**). Despite inflammatory activation, principal component analysis revealed that the mechanical properties of the hydrogel microenvironment remained a dominant determinant of macrophage polarization (**Figure 3f**). PC1 (53.4% variance), driven primarily by CD11b, HLA-DR, CD86, and CD80, separated LPS/IFN-γ-activated from control macrophages. PC2 (27.3% variance), with highest loadings for CD64, CD14, CD45, CD80, and CD206, separated the two mechanical environments, indicating that stress-relaxation modulate macrophage behavior independently of inflammatory co-stimulation.

These findings demonstrate that distinct stress relaxation values might serve as a critical mechanical cue directing macrophage polarization, with low τ_2_ matrices inducing pro-inflammatory phenotypes and high τ_2_ matrices supporting the regulatory/anti-inflammatory phenotypes.

### ECM τ_2_ drives transcriptional changes in macrophages

To investigate how ECM τ_2_ modulates macrophages transcriptomic profile, we encapsulated primary human macrophages from three donors in alginate hydrogels with either low or high τ_2_ properties and performed single-cell RNA sequencing before encapsulation (d0) and after 24h. The 24h timepoint was chosen to capture early gene expression programs, complementing protein-level measurements by flow cytometry and cytokine profiling at 48h. Uniform manifold approximation and projection (UMAP) dimensional reduction revealed distinct clustering between the two mechanical environments (**Figure 4a**), suggesting that matrix viscoelasticity induces distinct transcriptional profiles.

**Figure 4:**
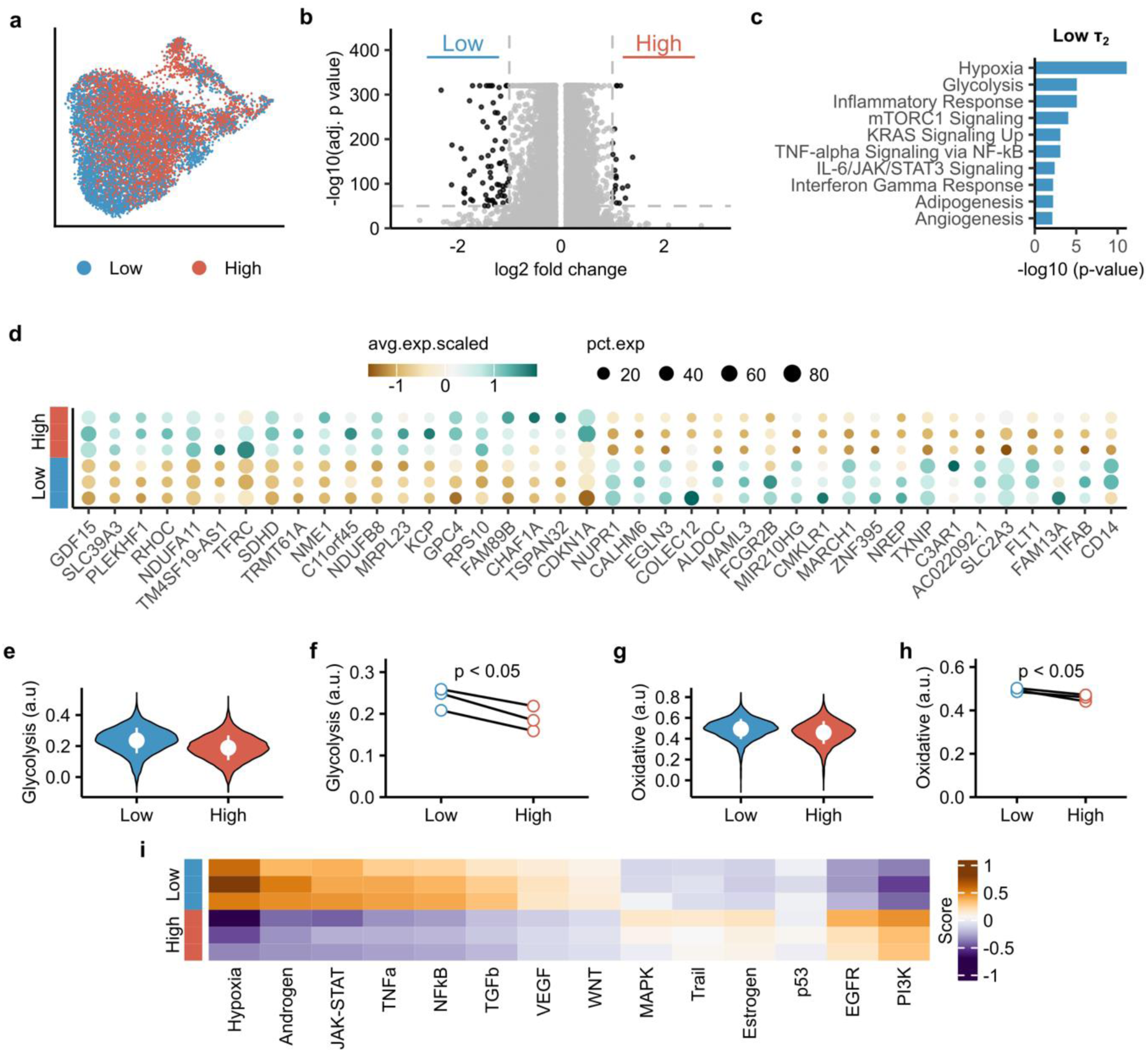
Low τ_2_ extracellular microenvironments activate pro-inflammatory, and high τ_2_ microenvironments anti-inflammatory pathway responses in macrophages. a) UMAP of scRNA-seq data from macrophages cultured for 24h in low or high τ_2_ alginate hydrogels reveals distinct spatial clustering based on hydrogel relaxation kinetics. Data are pooled from three donors. b) Volcano plots of differentially expressed genes (DEGs) comparing macrophages in low versus high τ_2_ conditions (pooled donors). Highly regulated genes were defined by a log₂ fold change > 1 and a p-value < 10⁻⁵⁰ and plotted in black, whereas dots in grey indicate log_2_ fold change < 1 and p-value > 10^−50^. c) EnrichR analysis using the MSigDB Hallmark 2020 database revealed enrichment of pro-inflammatory pathways among highly upregulated DEGs in low-τ₂ alginate hydrogels (pooled donors). d) Heatmap of the top 20 most regulated genes in low and high τ₂ matrices across three donors. The color scale reflects the average scaled expression, while the size of the dots represents the percentage of expressing cells within each condition. e) Single-cell KEGG pathway scoring for glycolysis (hsa00010) demonstrates elevated activity in low τ_2_ matrices. f) Mean pathway scores per donor confirm significantly higher glycolytic gene expression in low τ_2_ gels. g) Single-cell KEGG pathway scoring for oxidative phosphorylation (hsa00190) and h) corresponding mean expression per donor reveal slightly higher activity in low τ_2_ gels. i) PROGENy pathway activity analysis across three donors identifies hypoxia and PI3K signaling as the most differentially regulated pathways between low and high τ_2_ matrices.

For further downstream analysis we focused on highly differentially regulated genes (DEG) (|log₂FC| > 1, adj. p < 10⁻⁵⁰) (**Figure 4b**). Because of the large number of cells profiled (≈ 2000 cells per donor for both low and high τ_2_), statistical testing yielded extremely small p-values. This stringent cutoff was applied to highlight the most robust transcriptional changes and minimize false positives. Enrichment analysis using the MSigDB Hallmark 2020 gene set (via Enrichr) revealed significant activation of stress-and inflammation-associated pathways in the low τ_2_ gels, including hypoxia, glycolysis, and inflammatory response, as well as mTORC1, KRAS, and TNF-α signaling (**Figure 4c**). Hypoxia enrichment in the low τ_2_ gels was driven by canonical HIF downstream targets, including *SLC2A1, HK2*, *LDHA*, *VEGFA*, *PDK1*, and *NDRG1*, indicating activation of a hypoxia-responsive transcriptional signature rather than upstream oxygen-sensing regulators.

In contrast, enrichment analysis for macrophages in high τ_2_ gels yielded only two significant enriched terms (Myc Targets V1 and E2F Targets) under this strict cutoff, reflecting proliferative and biosynthetic regulation. When a more inclusive cutoff was applied (|log₂FC| > 0.5, adj. p < 10⁻³⁰), enrichment analyses recovered stress-and inflammation-associated programs in both, low and high τ_2_ gels, including TNF-α, mTORC1, and p53 signaling, but with additional metabolic features in high τ_2_ gels such as oxidative phosphorylation and adipogenesis (**SupplementaryFigure. 10**). Although macrophages from both low and high τ_2_ gels enriched TNF-α signaling via NF-κB, the contributing genes differed markedly between conditions. In the low τ_2_ gels, the top regulated genes included the pro-inflammatory cytokine *IL1B*, early inflammatory response transcription factors *FOS* (*AP-1*)^31^ and *CEBPD* (*C/EBPδ*)^32^, and upstream sensing components *TLR2* and *IFNGR2*. By contrast, macrophages in the high τ_2_ gels showed higher expression of genes linked to the noncanonical NF-κB arm (*NFKB2*)^33^, negative-feedback/attenuation (*DUSP2*^34^, *SPSB1*^35^), lipid-mediated regulatory programs (*SPHK1*^36–38^), and markers associated with tissue-resident/regulatory macrophage states (*EGR2*^39, 40^, *CSF1*^41^). A complete list of genes contributing to each enrichment term is provided in **Supplementary Table 4** and **Supplementary Table 5**. Although both low and high τ_2_ matrices activate stress-and inflammation-associated programs, macrophages in low τ_2_ gels preferentially show a bias toward acute pro-inflammatory, canonical NF-κB modules, whereas macrophages in high τ_2_ gels show a bias toward non-canonical NF-κB, feedback, and regulatory signatures.

A focused view of the top 20 regulated genes per condition across all donors further emphasized the metabolic and phenotypical changes (**Figure 4d**). Macrophages in high τ_2_ gels upregulated *GDF15*^42^, *NDUFA11*^43^, *NDUFB8*^44^, *SDHD*^45^, and other genes linked to mitochondrial activity and stress resilience, suggesting engagement of oxidative metabolism and regenerative phenotype. Conversely, low τ_2_ gels induced expression of immune-modulatory genes such as *CALHM6*^46^, *TXNIP*^47^, and *EGLN3*^48^ playing a role in regulating NLRP3 inflammasome activation, and sensing of hypoxia and redox changes. Interestingly, *TXNIP* has also been implicated in regulating cell mechanics and membrane tension^49^, suggesting that its upregulation in low τ_2_ gels may reflect direct macrophage adaptation to distinct τ_2_ microenvironments.

Given the transcriptional signatures pointing to metabolic reprogramming, a hallmark of macrophage polarization, we next quantified pathway activity at single-cell resolution. Glycolysis-related gene expression was significantly elevated in low τ_2_ gels both across pooled cells from all donors (**Figure 4e**) and at the donor-averaged level (**Figure 4f**), indicating a shift toward a more glycolytic phenotype reminiscent of pro-inflammatory macrophages. We also observed increased oxidative phosphorylation scores in low τ_2_ environments (**Figure 4g,h**), although this effect was less pronounced.

Next, we applied PROGENy pathway inference to identify potential pathway involved in sensing and reacting to different τ_2_ matrices (**Figure 4i**). Macrophages in low τ_2_ gels showed elevated activity of hpoxia, JAK-STAT, TNF-α, and NF-κB pathways; signatures commonly associated with acute inflammatory activation^50, 51^. In contrast, macrophages in high τ_2_ gels showed higher EGFR and PI3K activity, consistent with survival signaling, anti-inflammatory signaling and lipid metabolism^34, 52–54^. Together, these results demonstrate that matrix viscoelasticity serves as a potent regulator of macrophage polarization, coupling ECM stress relaxation to transcriptional profiles associated with inflammation, metabolism, stress adaptation, and immune function.

### Low τ_2_ primes macrophages for enhanced inflammatory activation independent of integrin-mediated cell-matrix adhesion

To further investigate how matrix stress relaxation influences macrophage polarization, we focused on the expression of *CD14*. CD14 emerged as one of the top markers in the loading of PC2 (**Figure 3**f). CD14 plays a critical role as a co-receptor for LPS recognition and is a hallmark of pro-inflammatory macrophages^55–57^. Macrophages encapsulated in low τ_2_ gels exhibited significantly elevated *CD14* expression compared to those in high τ_2_ gels (**Figure 5a**). Expression values are shown relative to cells before encapsulation (d0), allowing visualization of whether overall expression increased or decreased compared to the 2D baseline condition. This transcriptional upregulation was also reflected at the protein level, with low τ_2_ conditions exhibiting higher CD14 surface expression as measured by flow cytometry (**Figure 5b**).

**Figure 5:**
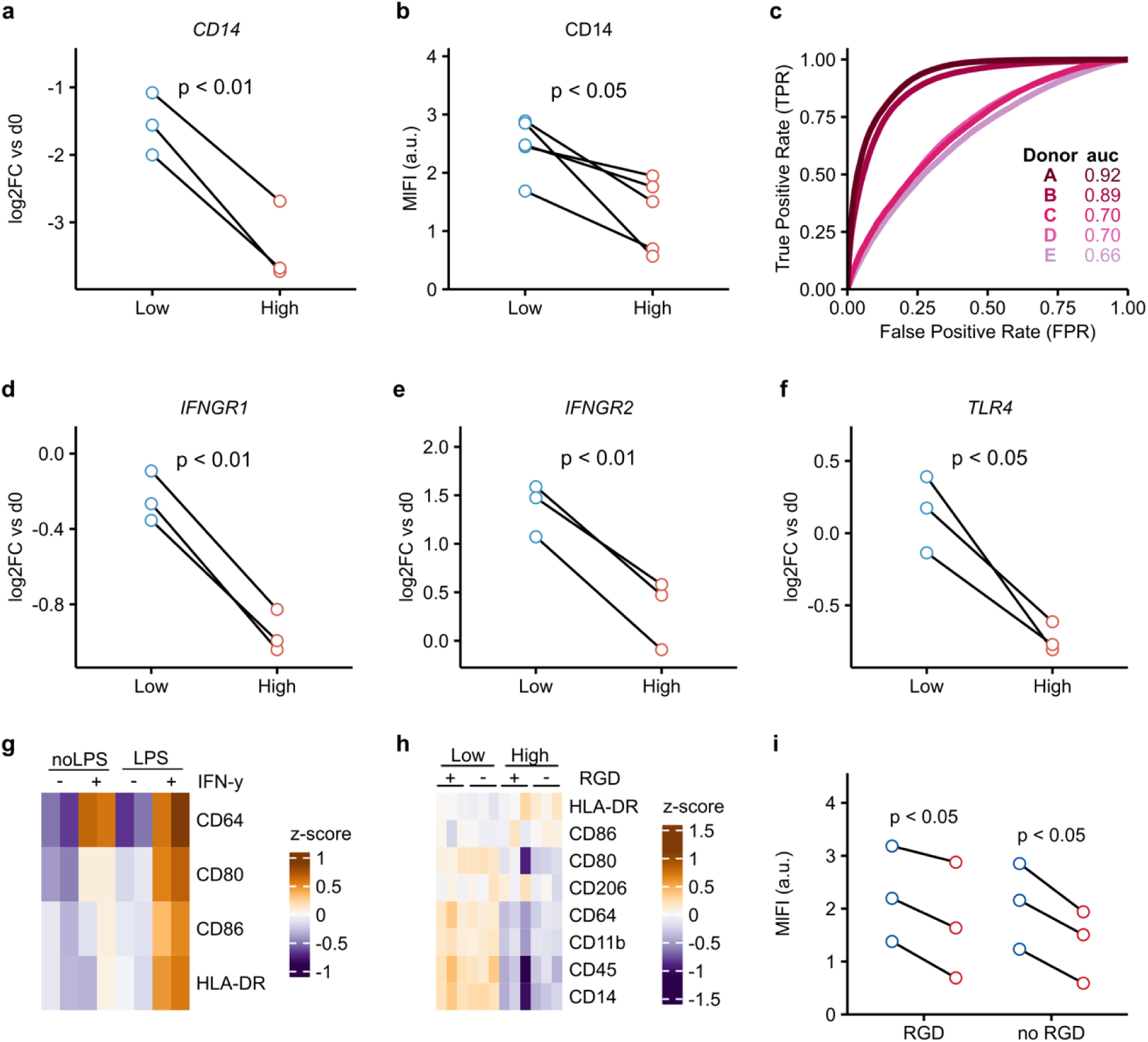
Low τ_2_ ECM stress relaxation enhances macrophage priming toward classical pro-inflammatory cues independently of cell adhesion. (a–b) CD14 expression is higher in macrophages cultured in low τ_2_ compared to high τ_2_ alginate hydrogels, as shown by **a)** pseudobulked mean RNA log2 fold change (log2FC vs. day 0, n = 3 donors) and **b)** CD14 protein levels measured by median logicle transformed fluorescence intensity (MlFI, n = 5 donors). **c)** Receiver operating characteristic (ROC) analysis of surface marker expression (CD11b, CD206, CD45, CD64, CD80, CD86, HLA-DR) revealed that CD14 exhibits donor-specific classifier performance in distinguishing macrophage activation states across low-and high-τ₂ matrices (n = 5 donors). **(d–f)** Expression of interferon-γ signaling receptors IFNGR1(d) and IFNGR2 (e) and LPS co-receptor TLR4 (f) is significantly upregulated in macrophages in low τ_2_ matrices relative to high τ_2_ ones (log2FC vs. day 0, pseudobulked RNA-seq, n = 3 donors). g) In 2D culture, combined stimulation with LPS and IFN-γ is necessary to induce broader pro-inflammatory surface marker expression (CD64, CD80, CD86, HLA-DR), whereas single-agent stimulation results in only partial or minimal activation (z-scored MlFI, n = 2 donors). h) Surface marker expression profiles of macrophages encapsulated in alginate gels with (+) or without (-) integrin-binding peptide RGD confirm that τ_2_ impacts surface marker expression in both adhesion-permissive and adhesion-deficient contexts (n = 3 donors). i) CD14 protein levels remain significantly elevated in low τ_2_ matrices regardless of the presence of RGD-mediated adhesion cues (MlFI, n = 5 donors). All statistical comparisons were performed using paired two-sided t-tests.

We next tested whether CD14 expression could predict the expression patterns of other surface markers associated with either low or high τ_2_ ECM. Receiver operating characteristic (ROC) analysis showed that CD14 expression could distinguish macrophages from low and high τ_2_ gels, although performance varied across donors (AUC = 0.67–0.92; **Figure 5c**). While classification was robust in some donors, lower AUC values in others indicate that CD14-based discrimination is not universally reliable and may be influenced by inter-individual variability.

To determine whether stress relaxing properties of the ECM could enhance responsiveness of macrophages to biochemical inflammatory signals, we analyzed the scRNA-seq data for key upstream pattern-recognition and cytokine receptors. Macrophages from low τ_2_ gels showed significantly increased expression of both interferon-γ receptor subunits, *IFNGR1* (**Figure 5d**) and *IFNGR2* (**Figure 5e**), as well as the LPS co-receptor *TLR4* (**Figure 5f**). Together with the increased expression of *CD14*, these findings suggest that macrophages in low τ_2_ ECM have heightened sensitivity to classical biochemical pro-inflammatory signals.

To test potential functional consequences of this receptor upregulation, we performed combinatorial stimulation assays in conventional 2D culture. As expected, CD64 responded robustly to IFN-γ alone, consistent with previous reports^58–60^. In contrast, optimal induction of the pro-inflammatory associated surface markers CD80, CD86, and HLA-DR required synergistic co-stimulation with both LPS and IFN-γ (**Figure 5g**). Neither stimulus alone was sufficient to achieve full activation, confirming the cooperative nature of classical macrophage polarization. Notably, CD14 expression remained unchanged in response to IFN-γ, either alone or in combination with LPS, indicating that its regulation under these conditions is not driven by canonical inflammatory stimuli but instead likely reflects a stress-relaxation dependent effect (**Supplementary Figure 11** and **Supplementary Figure 12**).

Next, we tested whether the influence of matrix viscoelasticity on macrophage polarization was mediated by cell adhesion. We generated alginate gels with or without the RGD adhesion motif. Regardless of RGD presence, macrophages encapsulated in low τ₂ gels upregulated CD11b, CD14, CD45, CD64, and CD80, whereas in high-τ₂ gels, CD86 and HLA-DR were upregulated (**Figure 5h**). Among these markers, CD14 showed a significant increase in expression in low τ₂ gels (**Figure 5i**). Since ROC analysis suggested CD14 as a predictor of the overall surface-marker response, these results indicate that the effects of matrix stress relaxation on macrophage polarization likely operate through adhesion-independent mechanisms.

### Macrophage transcriptomic profiles in low and high τ_2_ hydrogels recapitulate temporal progression of fracture healing *in vivo*

To test whether changes in ECM stress relaxation during fracture healing influence macrophage phenotypes *in vivo*, we compared RNA-seq profiles of macrophages cultured in low and high τ₂ gels with transcriptomes from macrophages isolated from early and late fracture hematoma phases. We used publicly available single-nucleus RNA-sequencing datasets from murine non-stabilized tibial fracture (GSE268276^61^, GSE234451^62^) and focused our analysis on days 3, 5, 7, because these had the most numbers of macrophages. We used DESeq2 to identify genes differentially expressed between macrophages cultured in low and high τ₂ gels and calculated the corresponding expression scores of these genes in macrophages from fracture hematoma.

UMAP analysis revealed distinct clustering of *in vivo* macrophages according to days post-injury. Module scores derived from genes upregulated in low and high τ₂ gels identified transcriptionally distinct macrophage populations (**Figure 6a, c**), indicating that gene programs associated with different matrix viscoelasticity correspond to discrete macrophage states *in vivo*. The low τ_2_ gene module exhibited the highest expression at 3 days post-injury and declined significantly over time (p = 2.8e-10; **Figure 6b**), consistent with the early pro-inflammatory phase switching towards anti-inflammatory phase. Moreover, the expression of receptors (*Tlr4*, *Ifngr1*, and *Ifngr2*) mediating biochemical pro-inflammatory macrophage activation by LPS and IFN-γ also declined progressively with increasing time post-injury (**Supplementary Fig. 9**). In contrast, high τ_2_ gene modules did not change significantly between days 3 and 7 (p = 0.25; **Figure 6d**). Together, these findings indicate that macrophage transcriptional responses to low τ_2_ matrices *in vitro* recapitulate the temporal gene expression profile of *in vivo* macrophages after bone fracture (**Figure 6e**).

**Figure 6:**
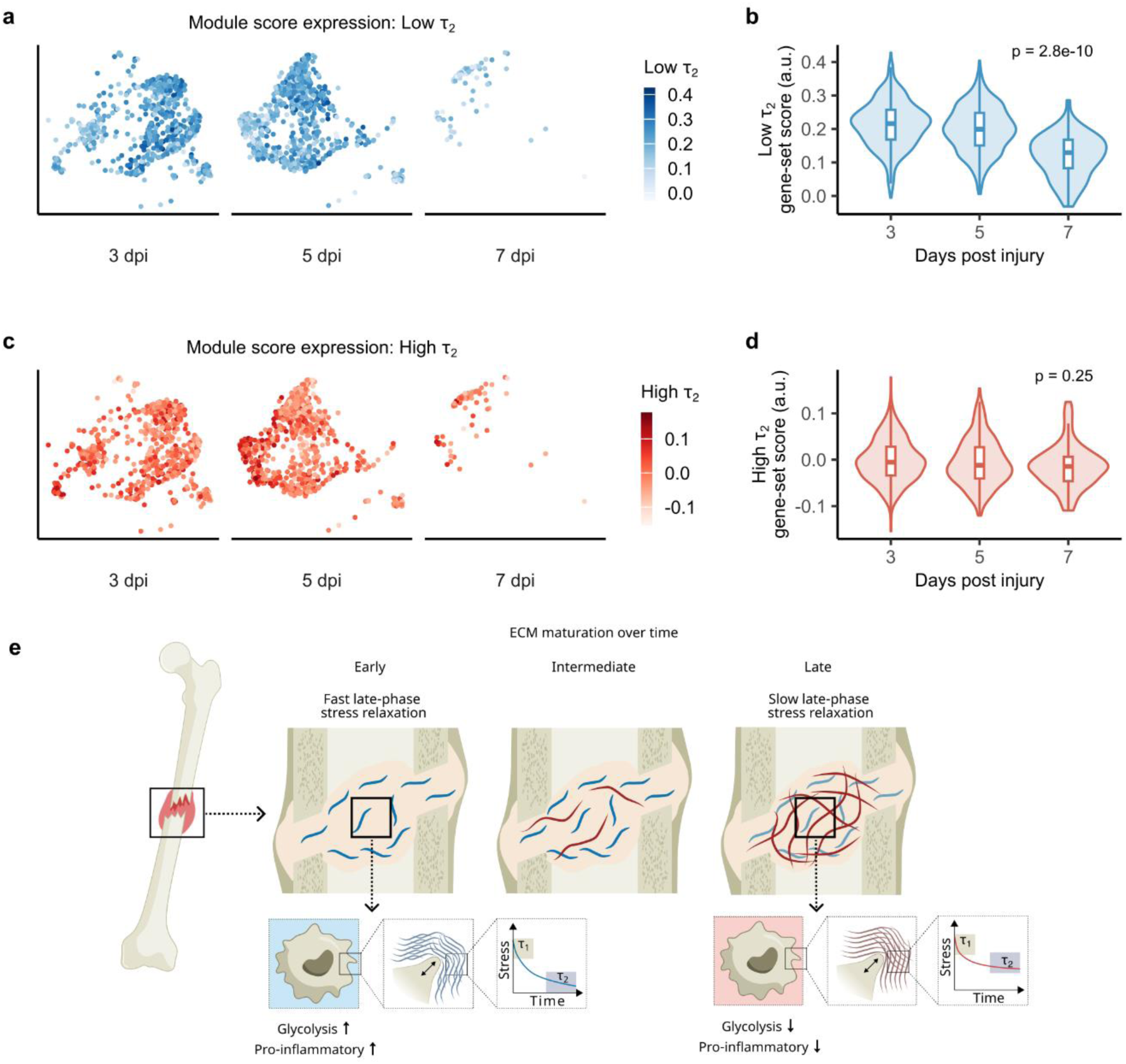
Transcriptional profiles of macrophages in hydrogels with low and high τ_2_ recapitulate gene expression profiles in early and late hematoma phases during bone fracture healing. (a,c) UMAP projections of macrophage transcriptomes from fracture hematoma at 3, 5, and 7 days post-injury, colored by module scores derived from **a)** low τ_2_ and **c)** high τ_2_ induced gene signatures. (b,d) Temporal progression of **b)** low τ_2_ and **d)** high τ_2_ gene module scores across 3, 5, and 7 days post-injury. Low τ_2_ signatures predominate early in healing and decline over time (p = 2.8e-10), while high low τ_2_ signatures did not change significantly over time. (p = 0.25). Individual points represent single cells, black line indicates mean trajectory. P-values calculated by Jonckheere-Terpstra test. **e)** Proposed model: Early post-fracture matrix exhibits low τ_2_ ECM, promoting glycolytic, pro-inflammatory macrophage phenotypes. As fracture hematoma mechanics mature toward higher τ_2_ ECM macrophages adopt less pro-and more anti-inflammatory/ tissue-remodeling phenotypes.

## Discussion

Here we reveal that distinct viscoelastic properties of the extracellular matrix steer macrophage polarization. Our mechanical characterizations of ex vivo human tissue samples revealed that ECM τ₂ stress relaxation properties positively correlated with time post injury during the hematoma phase, while no correlation was observed for ECM elastic parameters. Using an 3D *in vitro* cell culture system with tunable ECM mechanics to resemble differences found in ex vivo tissue, we could show that ECM τ₂ stress relaxation regulates macrophage polarization. Viscoelastic matrices with low τ_2_ steered macrophages into a pro-inflammatory phenotype, while viscoelastic matrices with high τ_2_ polarized macrophages into an anti-inflammatory phenotype.

Previous studies have demonstrated that macrophages sense mechanical properties through integrin-mediated adhesion and YAP/TAZ signaling in 2D environments. In contrast, our results indicate that in three-dimensional matrices τ_2_ modulate macrophage behavior through fundamentally different mechanisms. This could involve direct membrane deformation, cytoskeletal tension modulation, or mechanosensitive ion channel activation^16, 63^. The integrin-independent mechanosensing aligns with emerging evidence that macrophages in 3D environments possess expanded repertoires for detecting mechanical cues. For instance, macrophages in dense collagen hydrogels suppress pro-reparative gene expression (*FIZZ1*, *Ccl24*, *Fn1*) through cytoskeletal dynamics analogous to amoeboid migration, while monocytes in elastic versus viscous hydrogels upregulate F-actin independently of collagen adhesive ligands^20^.

Critically, the temporal scale of our identified mechanosensitive parameter is τ₂ = 500-2000 seconds. This temporal scale is markedly slower than the seconds-to-tens-of-seconds timescales characteristic of molecular clutch theory, where optimal cell spreading occurs when matrix stress relaxation times match integrin clutch binding and unbinding kinetics^64^. The difference in these timescales, combined with our observation that the effects are independent of integrin-mediated adhesion, indicates that alternative cellular dynamics operating on much slower timescales are governing adhesion independent macrophage mechanosensing in 3D. For instance, in nanoporous environments, cells exhibit distinct migration patterns depending on gel relaxation kinetics: fast-relaxing gels promote migration through long, shallow protrusions with nuclear positioning that functions as a mechanical piston, while slow-relaxing gels inhibit migration entirely^65^. This migration-based mechanosensing involves coordinated regulation by the actin cytoskeleton, microtubules, intermediate filaments, and mechanosensitive ion channels including TRPV4^66^. Filopodia dynamics are also modulated by the viscoelastic properties of the matrix. In fast-relaxing environments, filopodia exhibit average lifetimes of approximately 4 minutes, whereas in slow-relaxing matrices, their persistence extends to roughly 10 minutes^67^. These prolonged protrusions may facilitate mechanosensing by enabling cells to integrate mechanical information over extended timescales. Together, these observations support a model in which macrophages utilize a slower, adhesion independent mechanosensation in viscoelastic 3D environments that is distinct from the rapid, clutch-based sensing reported in 2D.

A crucial finding is that low τ_2_ matrices condition macrophages for enhanced inflammatory activation by upregulating gene expression of pattern recognition receptors (*TLR4*) and cytokine receptors (*IFNGR1*, *IFNGR2*). This mechanical conditioning of *TLR4* and *IFNGR1* creates potentially a state of heightened biochemical responsiveness that could amplify subsequent activation signals during tissue repair. This priming mechanism suggests that mechanical and biochemical signals operate synergistically rather than independently.

Although CD80 and CD86 are often co-regulated under inflammatory stimulation (e.g., LPS + IFN-γ), their regulation can diverge depending on context. CD80 is more tightly associated with classical pro-inflammatory polarization, whereas CD86 can also be upregulated by anti-inflammatory cytokines such as IL-4^68, 69^. Our observation that low τ_2_ matrices increased CD80 expression while high τ_2_ matrices increased CD86 and HLA-DR expression may therefore reflect distinct polarization trajectories; With low τ_2_ matrices steering macrophages to reach a more pro-inflammatory state, whereas high τ_2_ matrices bias them toward tissue-remodeling phenotypes.

The early dominance of low τ_2_ gene signatures followed by their progressive decline *in vivo* suggests that ECM network maturation, such as the resulting increase in matrix entanglement leading to higher τ_2_ values, reduces the pro-inflammatory expression profile of macrophages *in vivo*. For high τ_2_ gene signatures, however, our *in vivo* comparison did not reveal a corresponding increase with days post injury. This apparent discrepancy may be related to the use of a non-stabilized fracture model in the original publication of the dataset used here for the *in vivo* comparison. In non-stabilized fractures the switch from pro-to anti-inflammatory phase is delayed^70^. Accordingly, a substantially higher proportion of Arg1⁺ compared to CD206⁺ macrophages were still observed in the fracture callus at day 7 post-fracture^61^. Inclusion of later timepoints or the use of a stabilized fracture model would likely result in an increase of anti-inflammatory macrophages and, consequently, higher expression scores of high τ_2_ gene signatures.

Clinical implications could be derived from the notion that differences in ECM mechanics and formation of hematoma can (mis)guide crucial immune cell functions. ECM architecture, including collagen fiber orientation within the fracture gap, is known to determine healing outcomes^71^. Considering the decisive role of pro-and anti-inflammatory immune cell functions in bone healing^13^, aberrant hematoma ECM maturation (i.e., τ₂) during bone healing may serve as a predictive risk factor for delayed union or non-union outcomes. Further investigations into how dysregulated hematoma ECM mechanics contribute to impaired healing through aberrant immune regulation may provide novel insights into disease conditions that alter ECM properties and compromise healing capacity^72^. In vivo imaging modalities capable of assessing viscoelastic tissue properties, such as ultrasound^73^ or magnetic resonance^74, 75^ imaging, may ultimately enable screening for aberrant ECM remodeling during fracture healing and guide timely clinical intervention.

While our study provides evidence that matrix viscoelasticity regulates macrophage behavior, several limitations remain: First, we measured bulk mechanical properties, which may overlook local heterogeneity experienced by cells *in vivo*. High-resolution techniques such as atomic force microscopy or microrheology could map mechanical microenvironments at cellular scales. When combined with spatial transcriptomics, this could directly link local mechanics to immune cell states. Second, our hydrogels were mechanically static, limiting insights into how dynamic changes in τ₂ influence macrophage polarization. Future strategies incorporating degradable crosslinks or spatial gradients could enable both temporal and spatial tuning of viscoelasticity, providing tools to investigate how dynamic microenvironments regulate macrophage polarization^76–80^. Third, the relevance of these findings across other immune populations and tissue types remains to be determined. While prior work suggests that T-cells and monocytes also respond to viscoelasticity, systematic studies across immune contexts would broaden the translational impact.

This study reveals distinct viscoelastic properties of fracture hematoma as a fundamental regulator of macrophage polarization. Our findings support a model in which matrix viscoelasticity coordinates tissue repair by coupling macrophage polarization with viscoelastic ECM properties. Considering that hematoma formation and remodeling are hallmarks of regeneration after injury across many tissue types, ECM τ_2_ may hold regulatory roles beyond bone fracture. Our work highlights the important roles of ECM mechanical properties in immune cell regulation, with far-reaching implications for regeneration, chronic inflammation, cancer, and design of implants and biomaterials.

## Materials and methods

The collection of human tissue samples for this study was approved by the Ethics Committee of Charité – Universitätsmedizin Berlin (EA2/088/16).

### Material synthesis

Ultrapure sodium alginate (endotoxin level ≤ 100 EU/g) rich in guluronic acid blocks (>60%) was obtained from NovaMatrix in both high molecular weight (Pronova UP MVG, MW > 200 kDa, # 4200101) and low molecular weight (Pronova UP VLVG, MW < 75 kDa, # 4200501) forms. Alginate preparation was performed as previously described^81^. Briefly, RGD-alginate was synthesized by coupling the oligopeptide GGGGRGDSP (Peptide 2.0 Inc.) to the alginate backbone using carbodiimide chemistry. For the high-MW alginate, the reaction was carried out such that, on average, 20 RGD peptides were conjugated per alginate chain^82^. The molar peptide concentration was kept consistent for the low-MW alginate to ensure comparable functionalization. The modified alginate was then dialyzed (MWCO 3.5 kDa, CarlRoth, # 1978.1) against a decreasing salt gradient (150 mM to 0 mM NaCl) in deionized water, followed by sterile filtration (0.22 μm, Merck, # S2GPU05RE) and freeze-drying for long-term storage at –20°C. Freeze dried alginate was dissolved at 3% wt/vol in ImmunoCult^TM^-SF Macrophage Medium (StemCell, # 10961).

### Human monocyte isolation and macrophage differentiation

Primary human monocytes were isolated from fresh human blood from healthy donors by a density gradient with Lymphoprep (StemCell Technologies, # 18060) and SepMate-50 tubes (StemCell Technologies, # 85450) and negative isolation (EasySep Human Monocyte Isolation Kit, StemCell Technologies, # 19359) according to the manufacturers protocol. Monocytes were cultured in ImmunoCult^TM^-SF Macrophage Medium (StemCell, # 10961) at 1×10^6^ cells per ml with 1 % penicillin–streptomycin (ThermoFisher, # 15140) and 50 ng/ml recombinant human GM-CSF (BioLegend, # 572903) for 4 days.

### Stress relaxation measurements of human fracture hematoma

Human bone marrow hematoma was harvested during surgical stabilization of fractures 6 ± 3 d post-fracture and tissues were mechanically tested within 1 h after surgery. Collection of human fracture hematoma was approved by the institutional review board of the Charité – Universitätsmedizin Berlin (EA2/096/11). All patients gave their written consent. Hematoma samples were compressed at a rate of 1 mm min−1 until 15% strain, and then stress relaxation was measured over time for > 1000s.

### Uniaxial compression testing of alginate hydrogels

Cylindrical alginate gels (8 mm diameter, 3 mm height) were compressed to 15% strain at a rate of 0.025 mm/s using a BOSE TestBench LM1 system equipped with a 250 g load cell (Honeywell Model 31 Low). Stress was recorded for >2000 s while strain was held constant. The elastic modulus was calculated from the linear region between 8–12% strain. Stress relaxation data were low-pass filtered using a second-order Butterworth filter with a normalized cutoff frequency of 0.1 and then normalized to a maximum value of 1. The stress relaxation half-time was defined as the time point at which normalized stress decayed to 0.5.

### Stress relaxation modeling

Normalized stress relaxation data were fitted to generalized Maxwell models with 1 to n elements using non-linear least squares. Each Maxwell element consisted of an exponential decay term characterized by an amplitude (*A₁…Aₙ*) and a time constant (*τ₁…τₙ*). The fitting function followed the form:

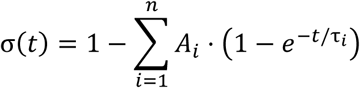

*σ(t)* denotes normalized stress, *A*_i_ the amplitude of the i-th element, and *τ_i_* its relaxation time constant. Model fitting was performed in R version 4.4.2 using the nlsLM() function from the minpack.lm (v1.2.4) package. Multiple initial parameter sets were generated per model, and the best fit was selected based on residual sum of squares. Parameter constraints ensured 0 < A_i_ < 1 and τ_i_ > 1. Model quality was evaluated using the coefficient of determination (R²), RMSE, Akaike information criterion (AIC), and Bayesian information criterion (BIC). Nested Maxwell models were formally compared using likelihood ratio tests. Pairwise comparisons between successively more complex models were performed by calculating chi-squared (χ²) statistic from the difference in log-likelihoods, usingthe corresponding difference in degrees of freedom.

### Encapsulation of macrophages in alginate hydrogels

Monocyte-derived macrophages were harvested after four days in culture using accutase (BioLegend, #423201), washed once with PBS, and resuspended in ImmunoCult^TM^-SF Macrophage medium at a concentration of 6.25 × 10⁶ cells/mL. Cell concentration was determined using a Countess 3 FL Automated Cell Counter (ThermoFisher). For hydrogel preparation, cell suspension was transferred into a Luer-lock syringe preloaded with a 3% (w/v) alginate solution, and gently mixed by repeatedly and carefully moving the plunger up and down. In a second syringe, 1.22 M calcium sulfate dihydrate (CaSO₄·2H₂O; ThermoFisher, # 315255000) was mixed in ImmunoCult^TM^-SF Macrophage medium to obtain a final calcium ion concentration of 27 mM for VLVG alginate gels and 15 mM for MVG alginate gels. The cell-alginate mixture and the CaSO₄-containing medium were connected using a female-female Luer-lock coupler and syringe mixed. The resulting mixture was immediately cast into custom-made cylindrical molds (8 mm diameter × 3 mm height) and allowed to crosslink for 30 minutes at room temperature. Gels were then transferred into individual wells of a 48-well plate containing 500 µL of ImmunoCult^TM^-SF Macrophage medium supplemented with 1% penicillin–streptomycin (ThermoFisher, #15140) and 50 ng/mL recombinant human GM-CSF (BioLegend, #572903). For pro-inflammatory polarization, the medium was further supplemented with 50 ng/mL IFN-γ and 100 ng/mL LPS. The final alginate concentration in all gels was 2% (w/v) and final cell density in the gels was 2×10^6^ cells /ml.

### Retrieval of cells from hydrogels

To retrieve cells from the alginate hydrogels, gels were transferred into a 12-well plate containing 2 mL of 20 mM EDTA solution (pH 7.4) per well. The plate was placed on an orbital shaker and incubated for 15 minutes, or until the gels were visually confirmed to be fully dissolved. The resulting cell suspension was collected into 10 mL tubes and washed with PBS prior to further downstream analyses.

### Multiplex ELISA assay

To assess cytokine secretion, cell culture supernatants were harvested from alginate-embedded macrophages. After 48h, culture medium from each well was collected and transferred into 0.5 mL protein LoBind tubes (Sigma-Aldrich, # 30108094). Samples were centrifuged at 1000 × g for 10 minutes at 4 °C to pellet residual debris, and the clarified supernatants were transferred to fresh tubes and stored at −80 °C until analysis. Cytokine levels were quantified using the LEGENDplex^TM^ Human Macrophage/Microglia Panel (BioLegend, # 740502) according to the manufacturer’s protocol. Samples were analyzed on a Sony ID7000 flow cytometer, and data were processed using LEGENDplex^TM^ Data Analysis Software Suite (BioLegend).

### Immunostaining and flow cytometry analysis

Following retrieval from hydrogels after 48h of culture in the gels, cells were stained using the Live/Dead Fixable Near-IR Dead Cell Stain Kit (ThermoFisher Scientific, #L23105) to assess viability. To prevent nonspecific (Fc receptor) mediated binding, cells were subsequently incubated with Human TruStain FcX^TM^ (BioLegend, #422302) and True-Stain Monocyte Blocker^TM^ (BioLegend, #426102). Surface marker staining was performed using a panel of fluorochrome-conjugated human-specific antibodies targeting CD45 (APC-FIRE750, BioLegend, #304062), CD11b (PE-Cy7, #101216), CD14 (AF488, #367130), CD206 (PE-Cy5, #321108), CD64 (BV421, #305020), CD80 (PE-Dazzle594, #305230), CD86 (BV650, #305428), and HLA-DR (AF700, #307626). Staining was conducted using Brilliant Stain Buffer (BD Biosciences, #566349). Samples were analysed on a SONY ID7000 spectral flow cytometer. Single-color controls were prepared using AbC Total Antibody Compensation Beads (ThermoFisher, #A10497) for antibody-based fluorophores and ArC Amine Reactive Compensation Beads (ThermoFisher, #A10346) for viability dye references. These controls were used to generate spectral reference libraries for compensation. Spectral unmixing was performed in the Sony ID7000 software using the Weighted Least Squares (WLS) method. Gating strategy was applied in FlowJo (v10.10.0), sequentially selecting cells, singlets, live cells, and CD45⁺CD11b⁺HLA-DR⁺ events. For downstream analysis, gated data were exported from FlowJo and processed in R (v4.4.2). Fluorescence intensities were transformed using the “logicle” transformation from the flowCore package (version 2.18.0) to accommodate the wide dynamic range and preserve negative values^83^. Median fluorescence intensities were calculated based on logicle-transformed data for statistical analyses and visualization. Principal component analysis (PCA) was performed on logicle-transformed and scaled data using the prcomp() function from the stats package (version 4.4.2) in R.

### Diffusion assay

Alginate hydrogels were incubated in 60 μg/mL GFP or 30 μg/mL tdTomato in a 96-well plate for 24 h at room temperature, protected from light. Gels were then rinsed once in PBS and transferred to a 48-well plate for imaging on a Keyence BZ-X810 epifluorescence microscope. Fluorescence images were batch-processed in FIJI/ImageJ v1.54g using a custom macro: each image was converted to 8-bit, segmented with Otsu thresholding, and area, mean intensity, standard deviation, and minimum intensity were measured. Results were exported as a CSV file and analyzed in R.

### Single cell RNA sequencing

Cells before encapsulation (0h) and after 24 h of culture in low and high τ_2_ hydrogels were labeled with human TotalSeq-C cell hashing antibodies (Biolegend) and were loaded into the 10x Genomics Chromium Controller according to the manufacturer’s protocol. Whole-transcriptome libraries were prepared using the 10x Genomics Single-Cell 5’ Library Preparation kit, and cell hashing libraries were prepared with the Chromium Single-Cell 5’ Feature Barcoding kit. cDNA was amplified for 12-14 cycles, yielding 14-150 ng/uL, libraries were sequenced on a NovaSeq X Plus (28 bp Read 1, 98 bp for Read 2) and raw reads were processed with CellRanger (v8.0.1), aligned to the human reference genome (GRCh38-3.0.0).

### Analysis of single-cell RNA-seq datasets

scRNA-seq data were processed and analyzed in R (v4.4.2) using the Seurat package (v5.2.1). Hashtag demultiplexing was performed using the function HTODemux() and positive.quantile = 0.99. Quality control was performed by filtering out cells with less than 200 detected genes or more than 10000 genes. Dimensionality reduction and integration were performed using Harmony, with donor specified as the integration variable. Highly variable features were selected using the “vst” method with 2,000 features, and confounding variables including total RNA count per cell (nCount_RNA) were regressed out. Cells were clustered at a resolution of 0.7. Analyses were focused on the 24 h timepoint to capture early transcriptional responses. Differential gene expression was calculated using Seurat’s FindMarkers function with the MAST statistical test, incorporating donor identity as a latent variable. Differentially expressed genes (DEGs) were filtered based on adjusted p-value < 10^−50^ and log_2_ fold change > 1, and pathway enrichment was conducted using enrichR (v3.4) against the MSigDB Hallmark 2020 gene sets (adjusted p-value < 0.05). To further quantify pathway-level activity, KEGGREST (v1.46.0) pathway scores were calculated for glycolysis (hsa00010) and oxidative phosphorylation (hsa00190). PROGENy (v1.17.3) was used to infer pathway activity scores from pseudobulk expression matrices.

### Analysis of published single-nuclei RNA-seq datasets

Publicly available single-nuclei RNA-seq datasets (GEO accession numbers: GSE268276 and GSE234451) were imported into R using the Seurat package (v5.2.1). Cells were filtered based on RNA content and mitochondrial and ribosomal transcript percentages: cells with fewer than 500 or more than 5,000 detected features, >5% mitochondrial transcripts, or >1.5% ribosomal transcripts were excluded. Filtered datasets were merged and normalized using the SCTransform workflow. Batch effects were corrected using Harmony integration. Muscle contamination (Ttn⁺ or Pax7⁺ cells) and low-quality clusters (low RNA content, low feature count, or high mitochondrial/ribosomal content) were removed. Cell types were annotated based on canonical marker expression and refined with SingleR using ImmGenData as reference. Macrophages were identified by co-expression of Ptprc and Adgre1; however, only 57 cells at day 0 and 11 cells at day 1 post-injury met this criterion, leading to exclusion of these timepoints from downstream analysis. Highly variable genes were identified using the VST method (2,000 features). Data were scaled while regressing out total RNA count per cell, mitochondrial transcript percentage, and ribosomal transcript percentage. Dimensionality reduction (PCA, UMAP) and graph-based clustering (FindNeighbors, FindClusters) were performed using the first 30 PCA dimensions, with a clustering resolution of 0.8. To evaluate the similarity between our *in vitro* single-cell RNA-sequencing (scRNA-seq) results and the *in vivo* datasets, a pseudobulk differential expression analysis was conducted on macrophages cultured in low and high τ_2_ hydrogels. Raw count matrices from the *in vitro* scRNA-seq data were aggregated at the sample level using Seurat’s AggregateExpression function, grouping by donor, low and high τ_2_, and timepoint. Aggregated expression data were subsequently analyzed using DESeq2 (v1.46.0) to identify differentially expressed genes (DEGs) between the high and low τ_2_ hydrogels. Genes with an adjusted p-value < 0.001 and an absolute log_2_ fold change > 0.25 were considered significantly differentially expressed. Human gene symbols from differential expression results were converted to their mouse orthologs using the R package orthogene (v1.12.0) using homologene as method and only one-to-one orthologs were retained. This gene set was then used to calculate a module score on the publicly available snRNA-seq datasets (GSE268276 and GSE234451). To assess directional trends in module score expression across experimental groups, the Jonckheere–Terpstra test was performed using the clinfun package (v1.1.5), and the associated p-value was reported.

### Statistical analysis

Statistical analyses were performed using R version 4.4.2. Comparisons between two groups were conducted using the unpaired/paired two-tailed Student’s t-test from rstatix package (v0.7.2). Correlation analyses were performed using Pearson’s correlation coefficient with the rcorr function from the Hmisc package version 5.2.3 in R. Linear regression analyses were performed using the lm function from stats package version 3.6.2 to assess relationships between variables. Assumptions of normality was evaluated using residual plots. P-values less than 0.05 were considered statistically significant unless otherwise noted. Error bars represent the standard error of the mean (SEM), unless otherwise specified. Receiver Operating Characteristic (ROC) curve analysis was performed using the pROC R package version 1.18.5 to evaluate the ability of CD14 expression to distinguish between low and high τ_2_ mechanical niches. The area under the curve (AUC) was reported as a measure of classification accuracy. All plots have been generated using ggplot2 (v3.5.2) and for illustrations inkscape (v1.3.2) was used. Heatmaps were created with ComplexHeatmap package (v2.22.0).

## Author contributions

R.S.K., D.J.M., K.S.B., and G.N.D. conceived the research. R.S.K. analyzed the human fracture hematoma dataset; developed the experimental setup; and performed and analyzed alginate hydrogel synthesis/modification, mechanical measurements, and *in vitro* experiments, including monocyte isolation, macrophage differentiation, flow-cytometric profiling, and cytokine assays. R.S.K., A.N., and D.M.M. performed the single-cell RNA-seq experiment, and R.S.K. analyzed the single-cell RNA-seq data. R.S.K., D.M.M., A.N., M.R.K., C.H.B., D.J.M., K.S.B., and G.N.D. interpreted the data. R.S.K. wrote the manuscript with input from all authors. All authors read and approved the final manuscript.

## Acknowledgements

We thank Simon Reinke (Core Unit “Cell Harvesting” of the Berlin-Brandenburg Centre for Regenerative Therapies) for providing human fracture hematoma samples and Dag Wulsten and Evi Lippens for biomechanical measurement of human fracture hematoma. This work was supported by the European Union (ERC-2021-ADG, 101054501), by the Einstein Foundation Berlin and by the German Research Foundation (Deutsche Forschungsgemeinschaft; SFB 1444)

