## Supplementary Figures for "Distinct matrix viscoelasticity in bone fracture hematoma steers macrophage polarization"

### Supplements

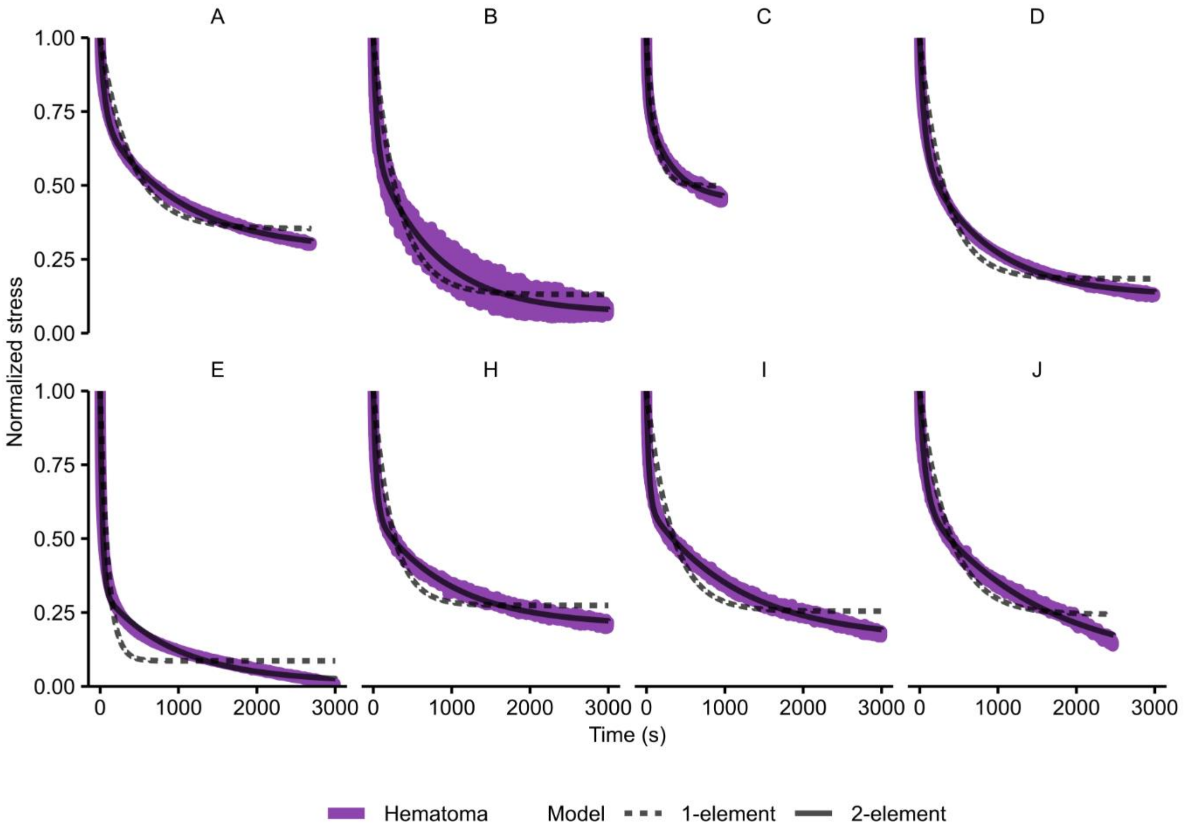


**Supplementary Figure 1: Stress relaxation behavior of fracture hematoma samples from individual donors and corresponding Maxwell model fits.** Normalized stress relaxation curves (purple) represent experimental data from hematoma samples of different patients (A–J; panel label indicates patient ID). Black dashed and solid lines show the corresponding one-element and two-element Maxwell model fits, respectively. Stress values were normalized to the initial peak stress.

**Supplementary Table 1:** ***Performance metrics of Maxwell models by number of elements (n).*** *Model performance is summarized across all fitted hematoma samples from one- to five-element Maxwell models. Metrics are reported as mean ± standard deviation (SD) and include the coefficient of determination ($R^2$), root mean square error (RMSE), Akaike information criterion (AIC), and Bayesian information criterion (BIC).*

| n | $R^2$ | RMSE | AIC | BIC |
|---|---|---|---|---|
| 1 | 0.82 ± 0.08 | 0.052 ± 0.01 | -1.4e+05 ± 3e+04 | -1.4e+05 ± 3e+04 |
| 2 | 0.99 ± 0.01 | 0.013 ± 0.007 | -2.7e+05 ± 8e+04 | -2.7e+05 ± 8e+04 |
| 3 | 0.99 ± 0.01 | 0.0081 ± 0.008 | -3.3e+05 ± 1e+05 | -3.3e+05 ± 1e+05 |
| 4 | 0.99 ± 0.01 | 0.0074 ± 0.009 | -3.6e+05 ± 1e+05 | -3.6e+05 ± 1e+05 |
| 5 | 0.99 ± 0.01 | 0.0073 ± 0.008 | -3.6e+05 ± 1e+05 | -3.6e+05 ± 1e+05 |

**Supplementary Table 2: Likelihood ratio tests for nested Maxwell models.** Pairwise likelihood ratio tests comparing one- through five-element Maxwell models. For each comparison, the $\chi^2$ statistic was calculated from the difference in log-likelihoods, with degrees of freedom equal to the difference in the number of parameters. The associated p-values indicate whether additional elements significantly improved the model fit.

| Patient | Model comparison | $\Delta\chi^2$ | p-value |
|---|---|---|---|
| A | n=1 vs n=2 | 1.6e+06 | 0 |
| A | n=2 vs n=3 | 2.5e+05 | 0 |
| A | n=3 vs n=4 | 2.5e+04 | 0 |
| A | n=4 vs n=5 | 9.4 | 0.0091 |
| B | n=1 vs n=2 | 1.6e+05 | 0 |
| B | n=2 vs n=3 | 4000 | 0 |
| B | n=3 vs n=4 | -0.00088 | 1 |
| B | n=4 vs n=5 | 85 | 3e-19 |
| C | n=1 vs n=2 | 2.3e+05 | 0 |
| C | n=2 vs n=3 | 1.6e+04 | 0 |
| C | n=3 vs n=4 | 2300 | 0 |
| C | n=4 vs n=5 | 50 | 1.6e-11 |
| D | n=1 vs n=2 | 1.5e+06 | 0 |
| D | n=2 vs n=3 | 4.3e+05 | 0 |
| D | n=3 vs n=4 | 4.8e+04 | 0 |
| D | n=4 vs n=5 | 1600 | 0 |
| E | n=1 vs n=2 | 9.2e+05 | 0 |
| E | n=2 vs n=3 | 5.8e+05 | 0 |
| E | n=3 vs n=4 | 2.6e+05 | 0 |
| E | n=4 vs n=5 | 3.3e+04 | 0 |
| H | n=1 vs n=2 | 1.2e+06 | 0 |
| H | n=2 vs n=3 | 1.1e+05 | 0 |
| H | n=3 vs n=4 | 6600 | 0 |

| Patient | Model comparison | $\Delta\chi^2$ | p-value |
|---|---|---|---|
| H | n=4 vs n=5 | 640 | 1.9e-139 |
| I | n=1 vs n=2 | 2e+06 | 0 |
| I | n=2 vs n=3 | 6.4e+04 | 0 |
| I | n=3 vs n=4 | 1.2e+04 | 0 |
| I | n=4 vs n=5 | 160 | 2.3e-36 |
| J | n=1 vs n=2 | 5.9e+05 | 0 |
| J | n=2 vs n=3 | 1.3e+05 | 0 |

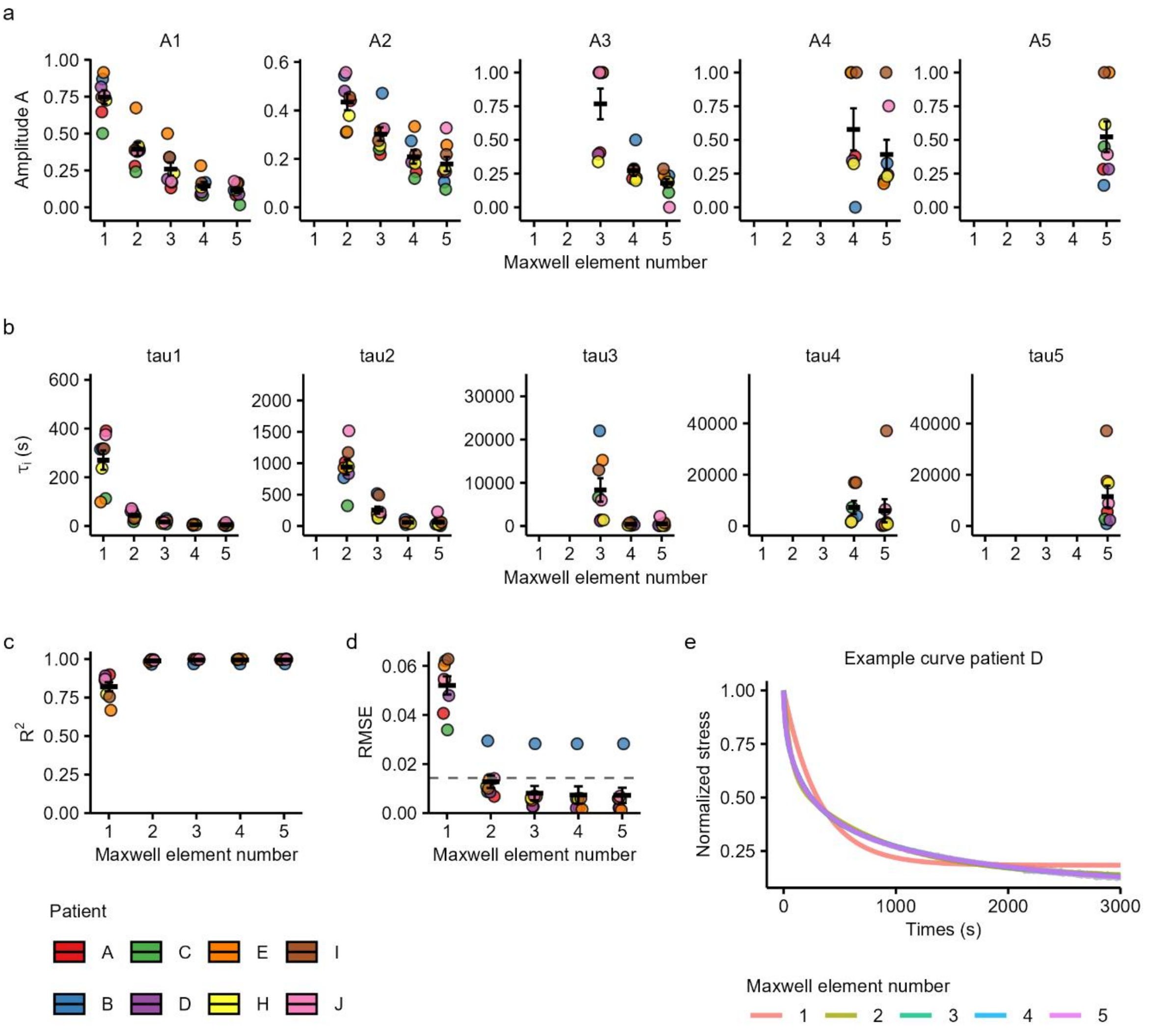


**Supplementary Figure 2: Evaluation of Maxwell model fits across patients and model complexity. a)** Fitted amplitude coefficients ($A_1$–$A_5$) from one- to five-element Maxwell models, reflecting the relative contribution of each viscoelastic element to the overall stress-relaxation response. **b)** Corresponding time constants ($\tau_1$–$\tau_5$) for each Maxwell element, describing the characteristic relaxation timescales in seconds. Both amplitude and time constant parameters were obtained by nonlinear least-squares fitting of experimental stress-relaxation curves from human fracture hematoma samples. Note: for four element fits, only for 7 out of 8 donors fit parameters were found. **c)** Coefficient of determination ($R^2$) showing goodness-of-fit for each model complexity; values approach 1 for higher-order models across all patients. **d)** Root mean square error (RMSE) of the fits. The dashed horizontal line indicates the estimated level of experimental noise. Noise was estimated by calculating, for each donor, the standard deviation of stress values during the final 10 seconds of the relaxation curve and then averaging these values across all donors. While RMSE decreases with increasing model order, the gain beyond the two-element model is marginal and falls within the noise threshold. **e)** Representative normalized stress-relaxation curves from a single patient (Patient D), overlaid with model fits for 1–5 Maxwell elements. Each dot represents an individual patient; colors denote patient identity (legend, bottom left), and lines represent mean ± SEM. The number of Maxwell elements used in the model is indicated on the x-axis of each subplot.

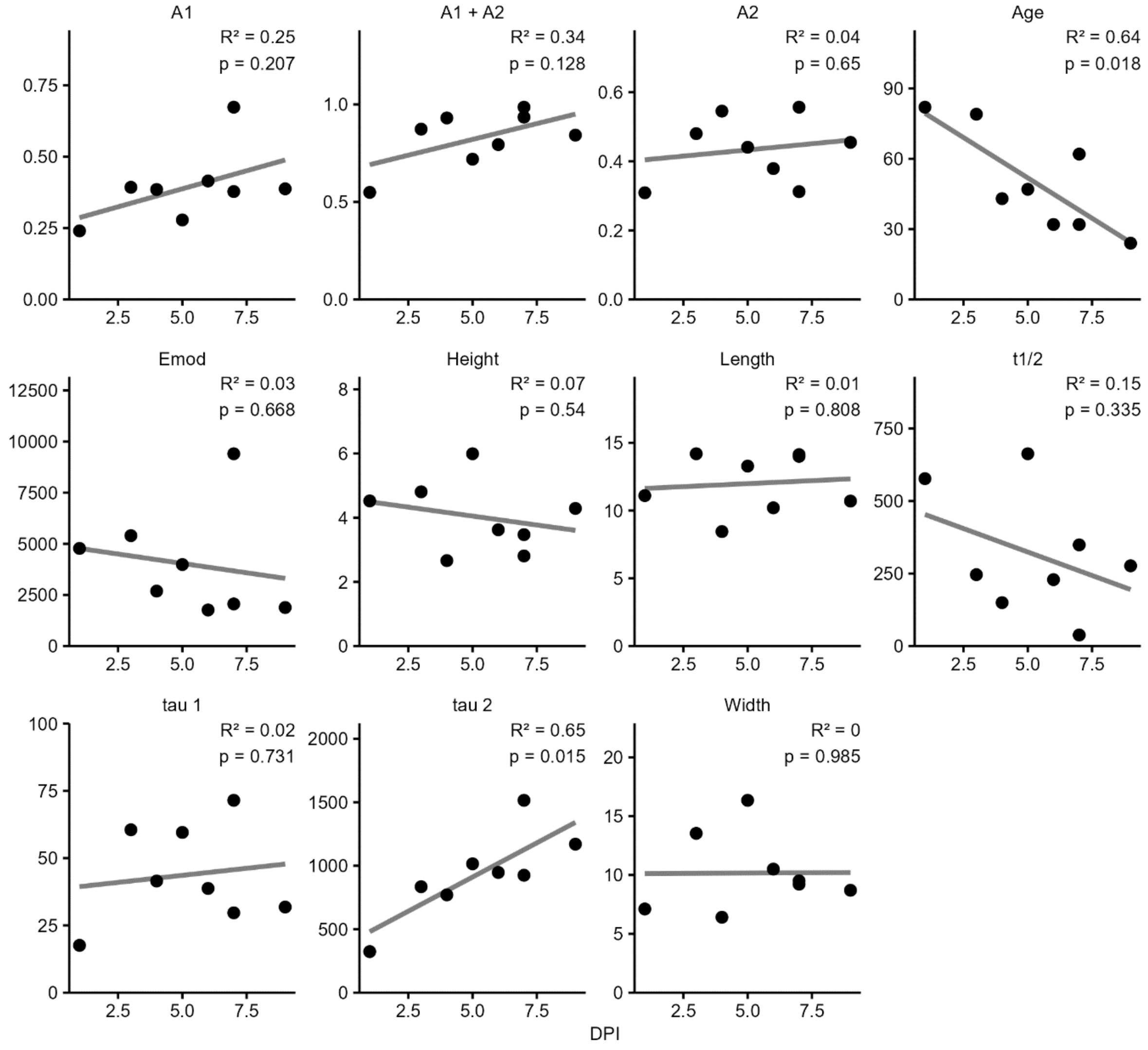


**Supplementary Figure 3: Pearson correlation of mechanical and sample parameters with days post-injury.** Linear regressions were performed to assess the relationship between days post-injury (DPI) and various mechanical and sample-derived parameters. Black dots indicate individual data points; gray lines represent linear fits. The coefficient of determination ($R^2$) and p-value are shown for each fit. A1, A2: amplitudes of Maxwell model elements, $t_{1/2}$: half-relaxation time, tau 1, tau 2: relaxation time constants, Emod: elastic modulus, height, width, and length indicate sample dimensions; age denotes patient age.

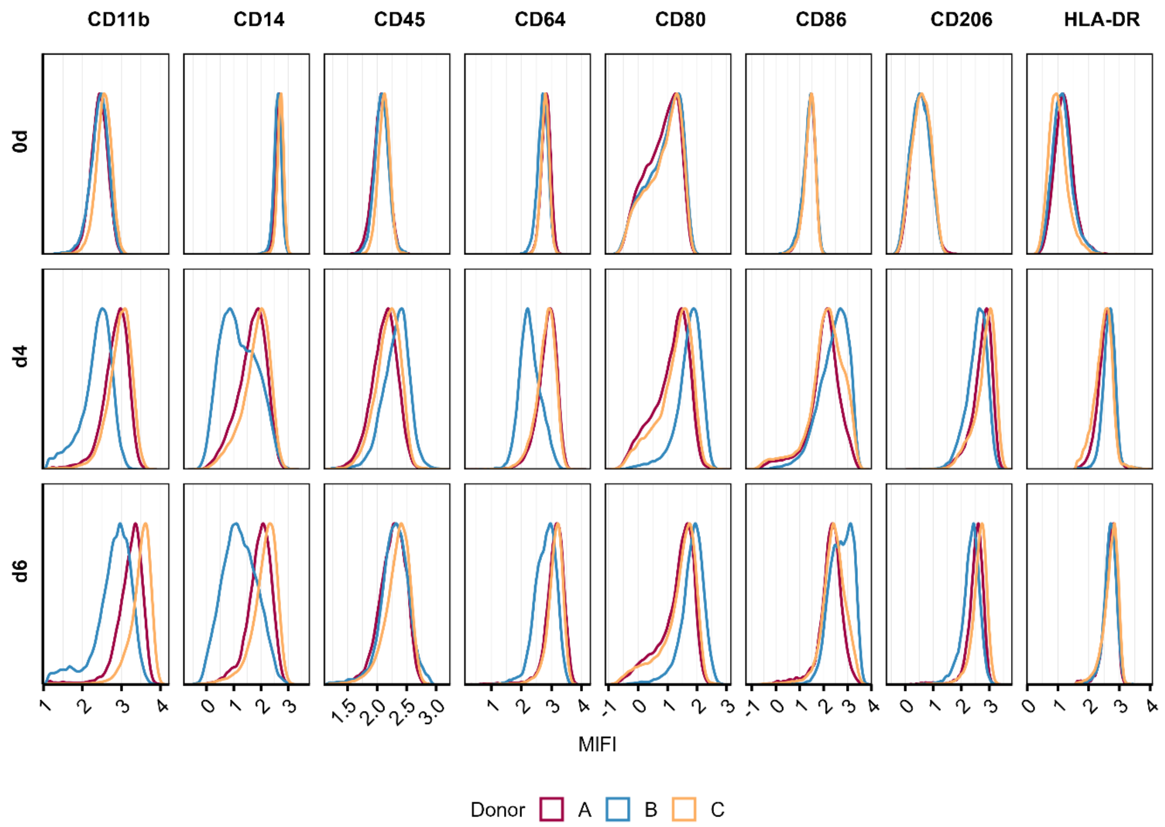


**Supplementary Figure 4: Expression of surface markers during GM-CSF–induced differentiation of monocytes.** Density plots show logicle-transformed mean fluorescence intensity (MIFI) of CD11b, CD14, CD45, CD64, CD80, CD86, CD206, and HLA-DR across three donors (A, B, C) at baseline (0d) and after 4 and 6 days of culture.

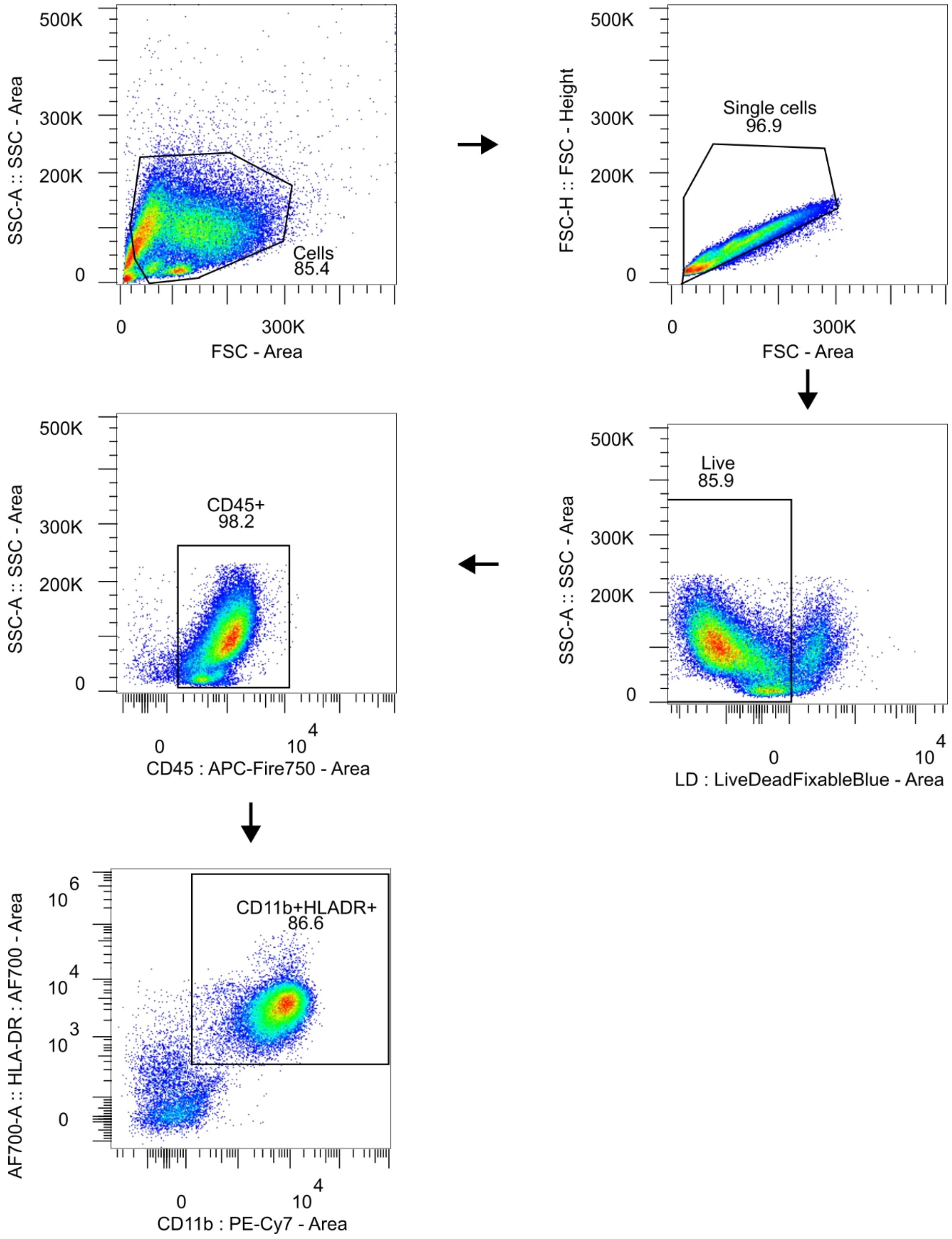


**Supplementary Figure 5: Gating strategy.** Example of monocytes stimulated with GMCSF for 4 days. Cells were sequentially gated on: Cells → Single cells → Live → CD45$^+$ → CD11b$^+$ HLA-DR$^+$.

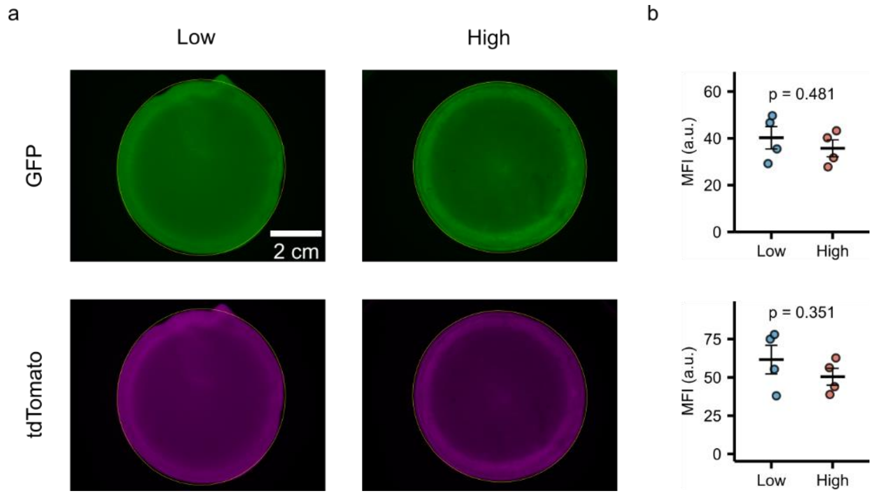


**Supplementary Figure 6: Diffusion of differently sized fluorescent proteins in low and high $\tau_2$ alginate hydrogels.** a) Representative images of gels incubated for 24h with GFP (27 kDa, top) or tdTomato (54 kDa, bottom) at calcium cross-linking concentrations: low: 27 mM and high: 15 mM. b) Quantification of mean fluorescence intensity (MFI) after incubation with GFP (top) and tdTomato (bottom). No significant differences were observed between low and high $\tau_2$ gels. n = 4 per condition, unpaired t-test.

**Supplementary Table 3: Summary of fitted parameters from the two-element Maxwell model applied to stress-relaxation data from low and high $\tau_2$ alginate hydrogels.** Reported values represent mean ± standard deviation (SD) across replicates (n=6).

| Alginate | tau1 | tau2 | A1 | A2 | A1 +A2 |
|---|---|---|---|---|---|
| Low | 28.89 ± 3.64 | 491.27 ± 109.97 | 0.58 ± 0.04 | 0.29 ± 0.04 | 0.87 ± 0.03 |
| High | 33.42 ± 4.29 | 1217.57 ± 167.17 | 0.21 ± 0.04 | 0.38 ± 0.03 | 0.59 ± 0.06 |

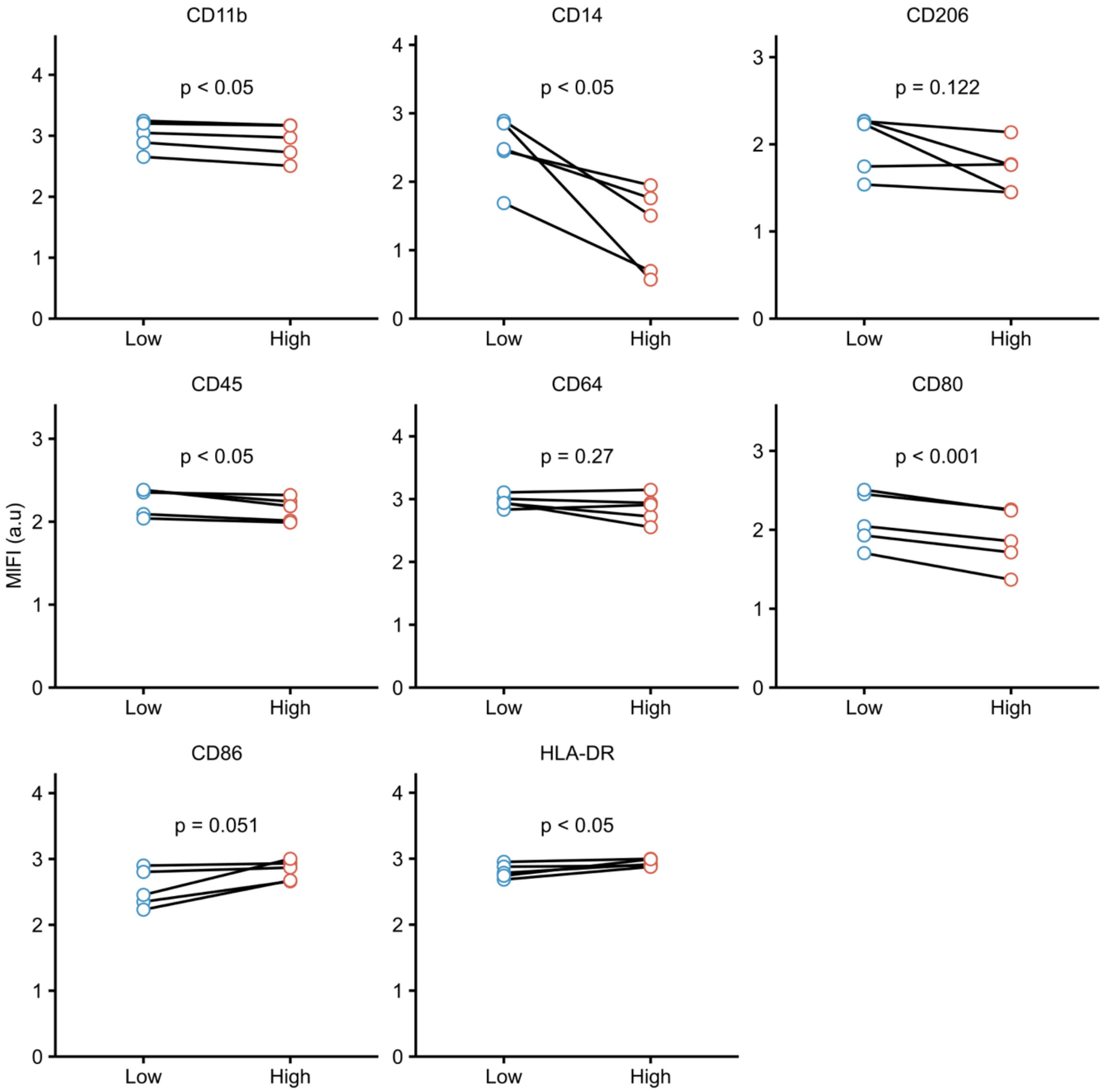


**Supplementary Figure 7: Surface marker expression in macrophages cultured in low and high $\tau_2$ alginate hydrogels.** Median logicle-transformed fluorescence intensity (MlFI) of indicated markers is shown after 48h of culture. Each line connects matched donors (n = 5). Statistical comparisons were performed using paired two-tailed t-tests.

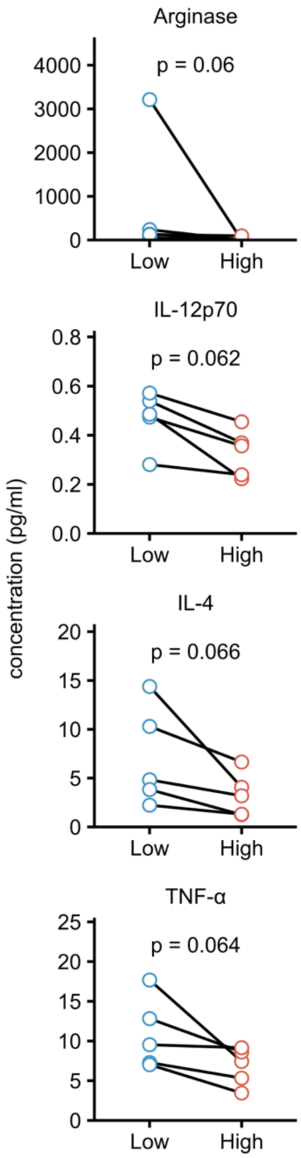


**Supplementary Figure 8: Cytokine secretion panel from macrophages in low and high $\tau_2$ alginate hydrogels.** Each line connects measurements from the same donor (n = 5). Statistical comparisons were performed using paired two-tailed t-tests.

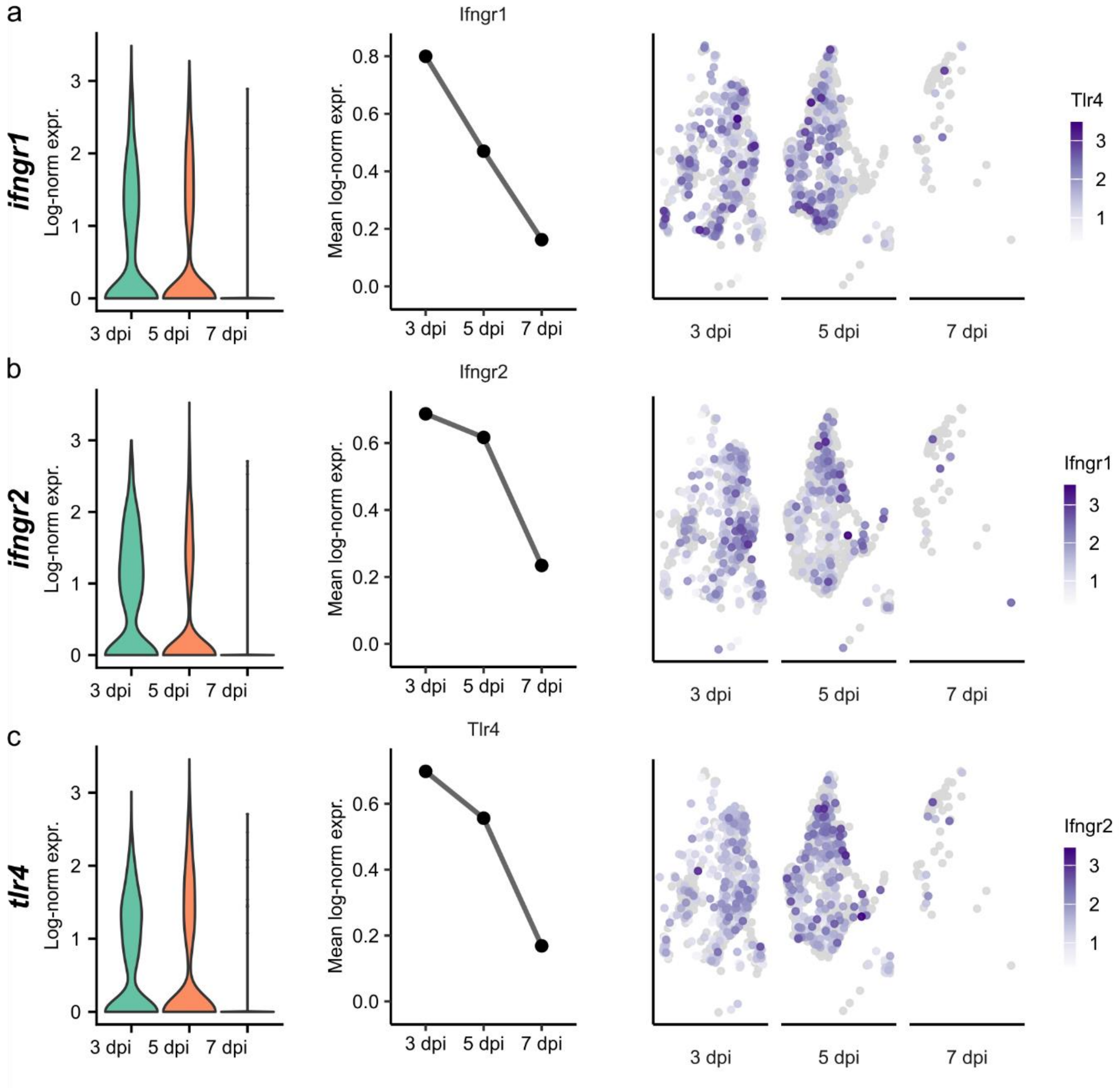


**Supplementary Fig. 9: Decline of ifngr1, ifngr2, and tlr4 with increasing days post-injury (dpi).** Single-nucleus RNA-seq data from GSE268276 and GSE234451 are shown for (a) *Ifngr1*, (b) *Ifngr2*, and (c) *Tlr4*. Left: violin plots of log-normalized gene expression across 3, 5, and 7 dpi. Middle: mean log-normalized expression levels across cells and timepoints. Right: UMAP visualization of expression patterns, colored by expression level.

r

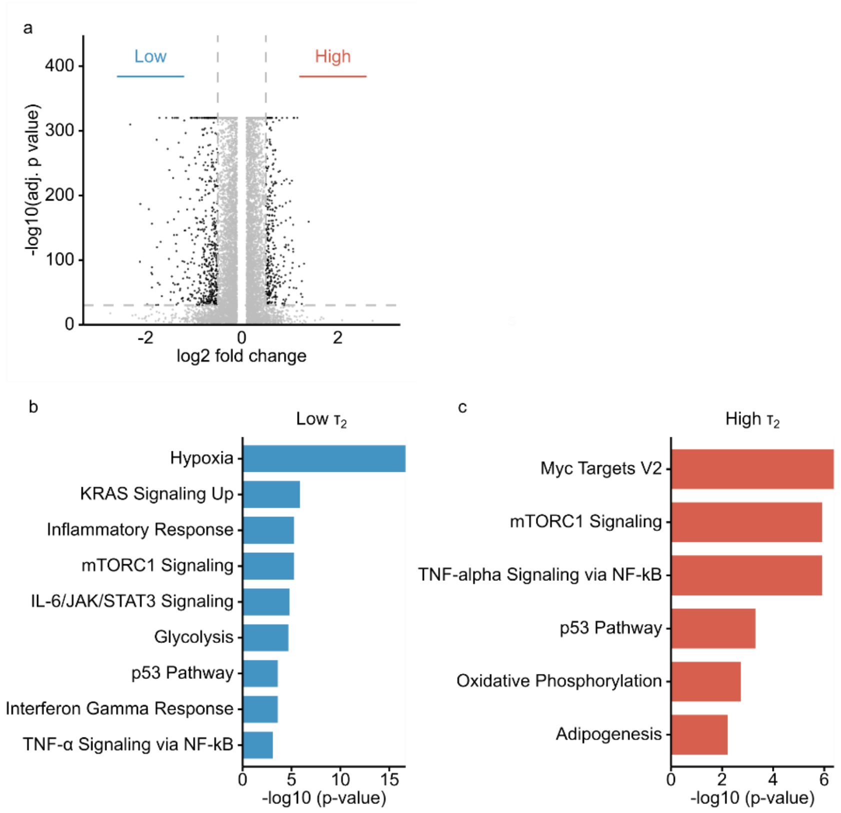


**Supplementary Figure. 10: Differential expression and pathway enrichment in low and high $\tau_2$ alginate hydrogels .** (a) Volcano plot of DEGs with cut-off values of $|\log_2 FC| > 0.5$, adj. $p < 10^{-30}$. (b) macrophages from low $\tau_2$ matrices show strong enrichment of hypoxia, glycolysis, and inflammatory pathways. (c) macrophages from high $\tau_2$ matrices show overlap with TNF-alpha signaling, but also proliferative signatures and oxidative metabolism.

**Supplementary Table 4: Enriched MSigDB Hallmark 2020 terms and associated genes identified by EnrichR in high $\tau_2$ alginate hydrogels.**

| Term | Genes |
|---|---|
| Myc Targets V2 | DUSP2;NOP16;GRWD1;WDR74;IMP4;NOC4L;FARSA;RRP9;BYSL |
| TNF-alpha Signaling via NF-kB | MAP2K3;EGR2;CDKN1A;DUSP2;CSF1;SPHK1;RHOB;NFKB2;SPSB1;SNN;PLAU;KLF9;TNFSF9;PLPP3;MSC |
| mTORC1 Signaling | MAP2K3;CDKN1A;TOMM40;TFRC;PNO1;TXNRD1;GSR;SLC1A4;RRP9;RIT1;PNP;HMBS;FDXR;CD9;EEF1E1 |
| p53 Pathway | PLK3;CEBPA;CDKN1A;NOTCH1;ABHD4;SPHK1;FDXR;OSGIN1;TNFSF9;TOB1;PHLDA3 |
| Oxidative Phosphorylation | NDUFB8;MTRR;ALAS1;ATP5PO;MFN2;ATP6V1H;NDUFC2;MRPS30;SDHD;ACO2 |
| Adipogenesis | SLC5A6;DDT;ATP5PO;RMDN3;LPL;PIM3;ACO2;TOB1;MGLL |

**Supplementary Table 5: Enriched MSigDB Hallmark 2020 terms and associated genes identified by EnrichR in low $\tau_2$ alginate hydrogels.**

| Term | Genes |
|---|---|
| Hypoxia | ERO1A;KDM3A;ZNF292;GBE1;SLC2A1;NEDD4L;ADM;SLC2A3;ENO2;RBPJ;SLC2A5;NDRG1;HK2;GYS1;MT2A;LDHA;ZFP36;MXI1;HMOX1;PDK1;CA12;BNIP3L;MAP3K1;MIF;FOS;VEGFA;SAP30;P4HA1;CCNG2;DDIT4;ALDOC;ALDOA;TMEM45A |
| KRAS Signaling Up | ERO1A;MAP3K1;JUP;IKZF1;CROT;RETN;TNFRSF1B;CSF2RA;DUSP6;ETV5;PTBP2;HSD11B1;LAT2;CLEC4A;IL1B;EPB41L3;C3AR1;ADAM8;CMKLR1 |
| mTORC1 Signaling | ERO1A;EGLN3;GBE1;SLC2A1;SLA;SLC2A3;TMEM97;IFI30;ADD3;HK2;DHFR;LDHA;P4HA1;SCD;DDIT4;NUPR1;ALDOA;PDK1 |
| Inflammatory Response | ABCA1;SEMA4D;IL1R1;IFNGR2;FPR1;ADM;SGMS2;TNFRSF1B;ACVR1B;ACVR2A;RNF144B;IL1B;CLEC5A;C3AR1;OLR1;CD14;CMKLR1;TLR2 |
| IL-6/JAK/STAT3 Signaling | ACVRL1;IL1R1;IL10RB;IFNGR2;IL1B;HMOX1;CD14;TNFRSF1B;ACVR1B;CSF2RA;TLR2 |
| Glycolysis | ERO1A;EGLN3;ZNF292;MPI;MIF;ENO2;HK2;VEGFA;SAP30;GYS1;LDHA;P4HA1;STMN1;DDIT4;MXI1;ALDOA;DSC2 |
| Interferon Gamma Response | MX2;FPR1;LYSMD2;IFI44;IFI30;SOD2;IL18BP;MT2A;EPSTI1;ST8SIA4;TXNIP;XAF1;FCGR1A;RAPGEF6;CMKLR1 |
| p53 Pathway | CCP110;FOS;IFI30;ACVR1B;NDRG1;DEF6;PITPNC1;ZFP36L1;RXRA;DDIT4;TXNIP;HMOX1;TAX1BP3;ERCC5;NUPR1 |
| TNF-alpha Signaling via NF-kB | ABCA1;TNFAIP8;CEBPD;IFNGR2;SLC2A3;FOS;SOD2;VEGFA;ZFP36;MARCKS;IL1B;OLR1;PTX3;TLR2 |

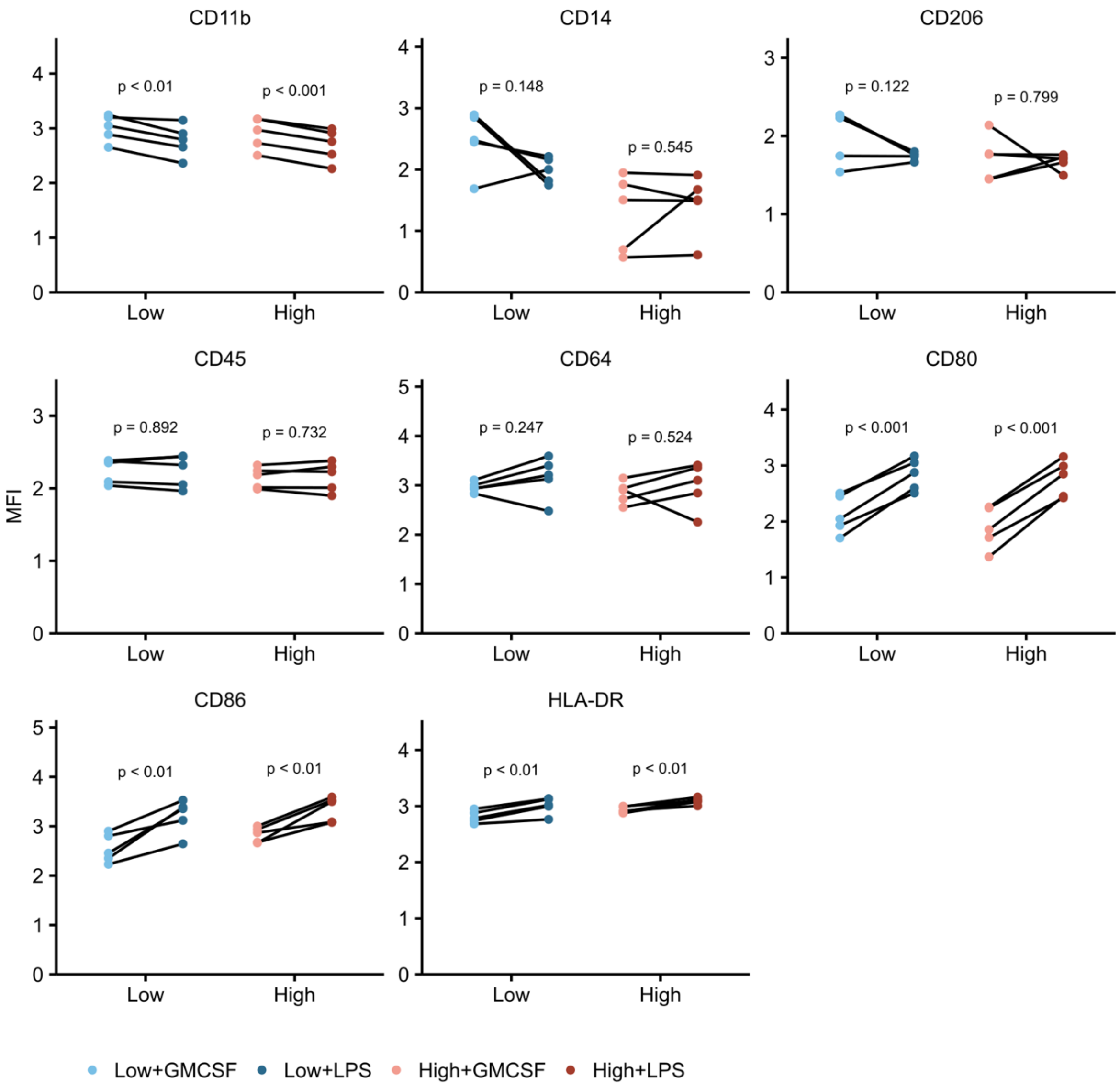


**Supplementary Figure 11: Upregulation of macrophage activation markers in low and high $\tau_2$ alginate hydrogels after LPS and IFN-γ stimulation.** Median logicle-transformed fluorescence intensity (MlFI) of surface marker expression in macrophages cultured for 48 hours in alginate hydrogels. Macrophages were either stimulated with GM-CSF (50 ng/mL; light blue and light red) or with LPS (100 ng/mL) and IFN-γ (50 ng/mL; dark blue and dark red). Lines connect matched donors. Paired two-tailed t-test, $n = 5$; * $p < 0.05$, ** $p < 0.01$, *** $p < 0.001$.

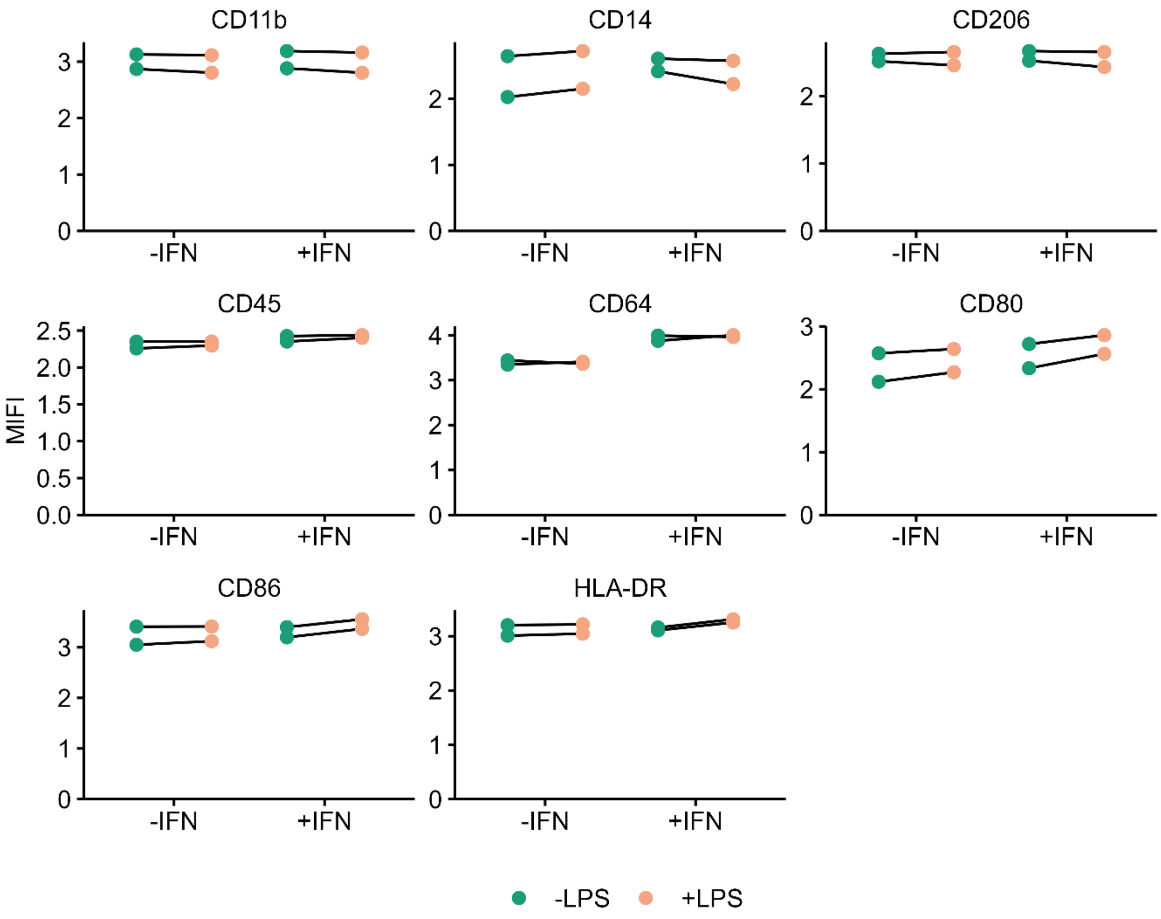


**Supplementary Figure 12: Expression of surface markers in macrophages following IFN-γ stimulation with or without LPS co-stimulation.** Macrophages were stimulated for 48h with IFN-γ alone (−LPS) or with IFN-γ and LPS (+LPS). Shown are median fluorescence intensities of logicle-transformed values (MIFI) for the indicated surface markers. Each line connects matched donors.
